# The rescue of epigenomic abnormalities in ICF1 patient iPSCs following *DNMT3B* correction is incomplete at a residual fraction of H3K4me3-enriched regions

**DOI:** 10.1101/2022.05.07.491011

**Authors:** Varsha Poondi Krishnan, Barbara Morone, Shir Toubiana, Monika Krzak, Maria Strazzullo, Claudia Angelini, Sara Selig, Maria R. Matarazzo

## Abstract

**Background:** Bi-allelic hypomorphic mutations in *DNMT3B* disrupt DNA methyltransferase activity and lead to Immunodeficiency, Centromeric instability, Facial anomalies syndrome, type 1 (ICF1). While several ICF1 phenotypes have been linked to abnormally hypomethylated repetitive regions, the unique genomic regions responsible for the remaining disease phenotypes remain largely uncharacterized. Here we explored two ICF1 patient-induced pluripotent stem cells (iPSCs) and their CRISPR/Cas9 corrected clones to determine whether gene correction can overcome DNA methylation defects and related/associated changes in the epigenome of non-repetitive regions.

**Results:** Hypomethylated regions throughout the genome are highly comparable between ICF1 iPSCs carrying different *DNMT3B* variants, and significantly overlap with those in ICF1-peripheral blood and lymphoblastoid cell lines. These regions include large CpG island domains, as well as promoters and enhancers of several lineage-specific genes, in particular immune-related, suggesting that they are pre-marked during early development. The gene corrected ICF1 iPSCs reveal that the majority of phenotype-related hypomethylated regions re-acquire normal DNA methylation levels following editing. However, at the most severely hypomethylated regions in ICF1 iPSCs, which also display the highest increased H3K4me3 levels and enrichment of CTCF-binding motifs, the epigenetic memory persisted, and hypomethylation was uncorrected.

**Conclusions:** Restoring the catalytic activity of DNMT3B rescues the majority of the aberrant ICF1 epigenome. However, a small fraction of the genome is resilient to this reversal, highlighting the challenge of reverting disease states that are due to genome-wide epigenetic perturbations. Uncovering the basis for the persistent epigenetic memory will promote the development of strategies to overcome this obstacle.

## BACKGROUND

In mammals, cytosine methylation occurs approximately at 70-80% of the CpG dinucleotides in the genome, and to a much lesser extent at non-CG sites [1, 2]. *De novo* establishment of DNA methylation occurs in embryos around the implantation stage by the combined action of the *de novo* DNA methyltransferases DNMT3A and DNMT3B, with a greater contribution of DNMT3B [3]. In mice the complete inactivation of these two enzymes leads to embryonic or postnatal lethality [4, 5], demonstrating their essential role in mammalian development.

In humans, biallelic hypomorphic mutations, mostly in the catalytic domain of DNMT3B, cause the Immunodeficiency, Centromeric instability, Facial anomalies type I (ICF1) syndrome. The main manifestation of this syndrome is a severe form of combined immunodeficiency that leads to lethal infections in early childhood [6, 7]. Additional clinical features are present in ICF syndrome patients at varying degrees of penetrance and severity and include intellectual disabilities, neurological defects, facial anomalies and developmental delay. While patients carrying *DNMT3B* variants represent the majority of ICF cases [6, 8–10], mutations in at least three additional genes can cause ICF syndrome, highlighting the complex genetic heterogeneity of the disease [11, 12]. In ICF1 patients, DNA hypomethylation is apparent at specific heterochromatic regions, including the pericentromeric and subtelomeric repeats [13, 14]. The ensuing defect in condensation of the pericentromeric regions of chromosomes 1, 9 and 16 contributes to the centromeric instability in immune cells, while subtelomeric hypomethylation leads to telomere shortening and premature senescence of patient fibroblasts [14, 15]. The humoral immune deficiency, characterized by low to absent levels of IgA, IgG and IgE, is a common clinical phenotype in ICF patients, even though it is slightly more pronounced in ICF1 compared to patients of the other subtypes [6]. Agammaglobulinemia in ICF patients has been attributed to disturbances in peripheral B cell maturation, as patient peripheral blood contains naive B cells, but lacks memory and plasma cells [16].

Despite the identification of ICF syndrome-causative genes, very little is known about the underlying molecular mechanisms and the cascade of pathophysiological events that lead to this severe clinical phenotype, as well as the other clinical phenotypes. Particularly, it is yet unknown which of the methylation defects throughout the genome are directly responsible for the aberrant gene expression pattern and the cellular and developmental defects typical of ICF syndrome. DNA methylation and transcriptional studies of ICF1 patient whole blood and lymphoblastoid cell lines (LCLs) identified several key alterations in epigenetic and gene expression profiles [17–21]. We previously demonstrated that pathogenic *DNMT3B* variants disrupt the intragenic methylation level and the silencing of alternative and cryptic promoters in several genes involved in the regulation of immune cell function, thereby interfering with the control of transcription initiation [20]. This is consistent with recent genome-wide studies describing that intragenic CG methylation of active genes is dependent on DNMT3B activity in both humans and mice [22–25]. Despite these important insights, it is difficult to distinguish between the direct genomic targets of DNMT3B during implantation, which constitute the primary methylation defects, from the secondary methylation defects that occur later in development. The latter most probably arise due to genome-wide perturbations of the epigenetic landscape induced by the primary DNA methylation changes.

Generation of ICF1 patient-derived induced pluripotent stem cells (ICF1 iPSCs) provide a powerful model for studying the earliest pathogenic events in ICF1 syndrome and for identifying the genomic regions prone to DNA methylation loss as a consequence of *DNMT3B* loss of function (LOF) variants [26, 27].

Indeed, ICF1 iPSCs retain the characteristic hypomethylation at pericentromeric and subtelomeric repeats [14, 26]. Consistently, when differentiated into fibroblast-like cells, they recapitulate the abnormal telomere shortening and premature senescence phenotypes, originally described in ICF1 patient fibroblasts [26]. Thus, although DNMT3B is highly active in human iPSCs, its low residual activity in ICF1 cells is sufficient to permit reprogramming and pluripotency, thereby providing a reliable platform to study the etiology of ICF1 syndrome and the dynamics of epigenetics abnormalities in the context of early development.

By performing homology directed repair (HDR) based gene-editing through the CRISPR/Cas9 technology we recently corrected *DNMT3B* mutations in ICF1 iPSC lines derived from two patients [28]. Focusing on repetitive regions in the corrected iPSCs, we revealed that while pericentromeric repeats acquired normal methylation levels, DNA methylation at the majority of subtelomeres was only partially restored.

Importantly, we found that the chromatin context at subtelomeres is significantly altered in both somatic and patient-derived iPSCs, thus inhibiting full DNA methylation rescue by the corrected DNMT3B.

Therefore, our ICF1 iPSCs, together with their corrected isogenic counterparts, offer a unique opportunity to decipher the molecular pathogenesis in ICF1 syndrome, as well as provide insights into the factors that regulate the tethering of DNMT3B to specific regions in the genome of pluripotent stem cells. Several studies have reported that CG content and chromatin context influence the recruitment of *de novo* DNMTs [22]. In addition, transcription factors with preferential binding to GC-rich sequences, may also shape DNA methylation patterns by recruiting DNMT enzymes [29, 30]. However, the exact role of these factors and their contribution to the targeting and activity of DNMTs at various genomic regions is not fully known.

Here, we examined ICF1 iPSCs and their corrected counterparts through a systematic and integrated multi-omic approach focusing on the non-repetitive portion of the genome. By comparing the ICF1 iPSCs with WT iPSCs, we identified key targets of *de novo* methylation by DNMT3B, and gained insights into its deficient activity at early disease stages and genes that are potentially involved in generating the abnormal ICF1 phenotypes. Following correction, we utilized the isogenic clones to identify regions which partially or fully restored normal DNA methylation patterns. These findings contribute to the identification of molecular factors that influence the ability to correct the ICF1 abnormal epigenome.

## RESULTS

### DNA hypomethylation patterns in different ICF1 iPSCs and their rescue in corresponding corrected clones are highly comparable

To extensively evaluate the genome-wide modified epigenetic landscape associated with DNMT3B LOF, we utilized iPSCs derived from pR and pG ICF1 patients and their respective CRISPR-corrected clones [26, 28]. The pR patient is homozygous for the D817G change in the DNMT3B catalytic domain, while the compound heterozygous pG patient carries the I41fsX42 mutation leading to a premature stop codon in one *DNMT3B* allele and a mutation leading to the S780L change in the catalytic domain of the second allele. Two isogenic clones of pR iPSCs, cR7 and cR35, have biallelic correction of the missense mutation. iPSC clones cG13 and cG50 clones have corrected the null mutation in pG iPSCs, and maintain the missense mutation in the catalytic domain of the protein, thus mimicking the genotype of heterozygous carriers that are healthy.

To investigate the effect of the pathogenic *DNMT3B* variants on DNA methylation levels across the genome, and to identify the regions rescued by the corrected DNMT3B, we performed whole genome bisulfite sequencing (WGBS) in WT, patient and corrected iPSCs. Consistent with the hypomorphic nature of the *DNMT3B* mutations in ICF1 pR and pG patients, we observed a decrease, albeit mild, of global methylated CG (mCG) levels (from 74.8% in WT-iPSCs to 70.1% in pR iPSCs and 64.6% in pG iPSCs) (**Fig. S1A)**. A decrease in non-CG methylation in pR and pG iPSCs was also detected (**Fig. S1B, C**), confirming previous observations in additional ICF1-derived iPSC lines [27]. Although the global mCG levels in patient iPSCs were not extensively compromised, we identified 36,876 and 31,854 hypomethylated regions in pR and pG, respectively, when we searched for differentially methylated regions (DMRs) between the two ICF1 iPSCs and our internal control WT1 iPSCs. To more accurately define the DMRs, we compared the methylation levels of WT1 iPSC to that of an additional WT iPSC from a public dataset, which we designated as WT2 iPSC (GSM1385983, derived from human dermal fibroblasts [31]). We then filtered out the hypo-DMRs whose methylation level differed between WT1 and WT2 (difference between the ratio values > 0.2). Following this step, we retained 26,031 and 22,727 hypo-DMRs for pR and pG respectively. A significant fraction of these hypo-DMRs strictly overlapped between the pR and pG samples (n=8560, p-value < 0.0001; shuffle test), and the majority (62%) were in proximity (within 20kb), supporting their relevance in the context of disease pathogenesis **(Fig. S1D)**.

We next clustered the hypo-DMRs into four groups based on their DNA methylation levels in patient iPSCs, with Groups 1 and 2 including the regions with the lowest values compared to the WT1 and WT2 controls (**Fig. 1A**). Overall, we observed that the ability to re-acquire WT methylation levels following *DNMT3B* correction was inversely proportional to the degree of hypomethylation in the original ICF1 iPSCs. Hence, while the less hypomethylated regions in pR and pG iPSCs (Groups 3 and 4) regained almost full methylation in the corrected clones, the recovery at regions with severe hypomethylation (Groups 1 and 2) was highly compromised (**Fig. 1A**).

**Figure 1.**
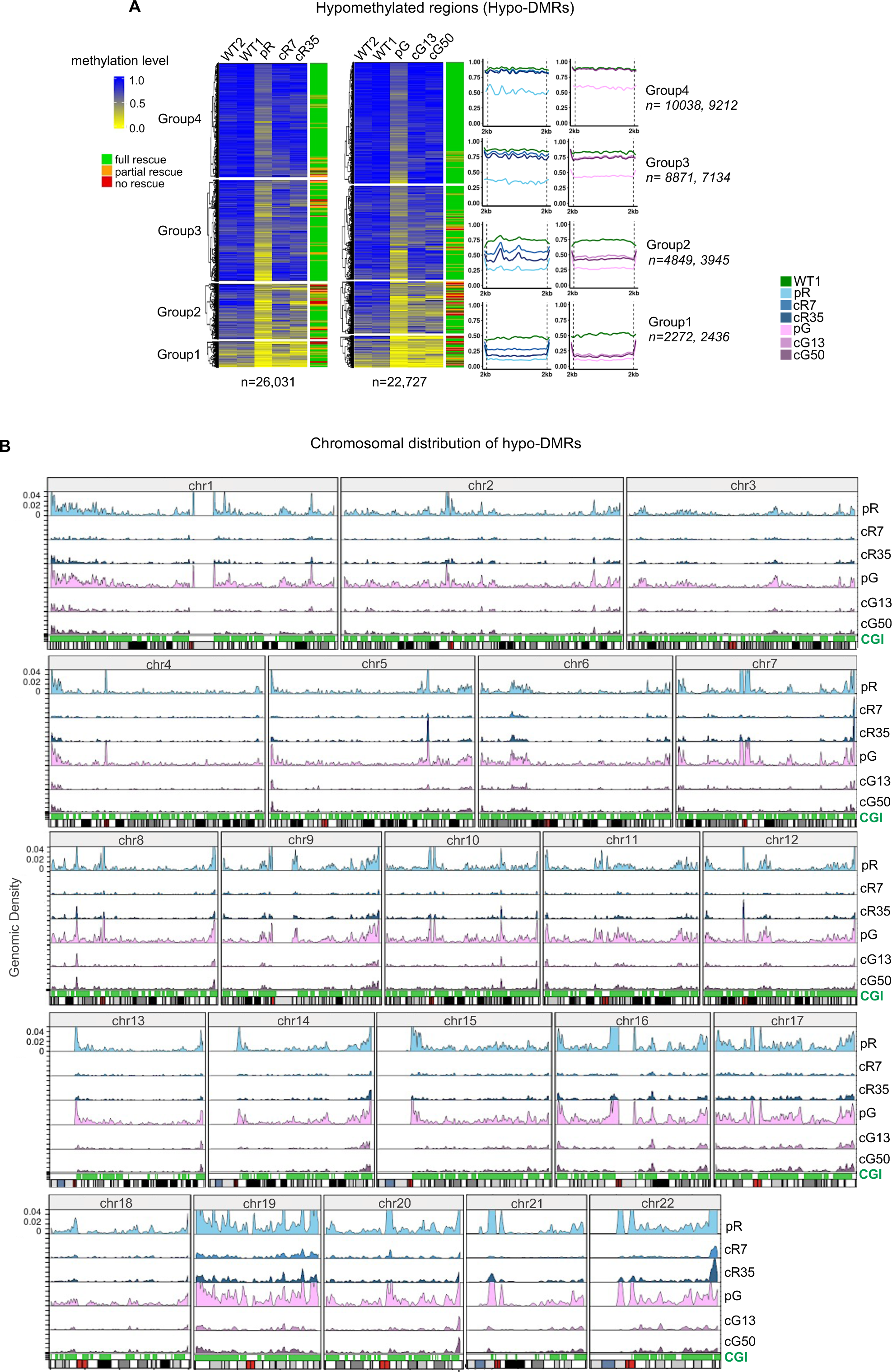
DNA hypomethylation profiles are conserved between different ICF1 iPSCs. **(A)** Heatmap representation of methylation levels as determined by WGBS (expressed as ratio of the number of Cs over the total number of Cs and Ts) at hypo-DMRs of size 1kb in pR and pG ICF1 iPSCs, their respective corrected clones and controls WT1 and WT2. The hypo-DMRs were clustered using k-means into four groups based on the methylation level across ICF1 iPSCs. The last column indicates the ICF1 hypo-DMR rescue status: full, partial and no rescue of ICF1 hypo-DMRs indicate remethylation in both, single, or none of respective corrected clones. The profile plots on the right depict the average methylation level of each subgroup of hypo-DMRs and their flanking regions (±2kb). The numbers of hypo-DMRs in pR and pG iPSCs in each group are indicated on the right. **(B)** Chromosomal distribution of hypo-DMRs in pR and pG iPSCs and those persisting in the respective corrected clones (cR7=3,599, cR35=5,171; cG13=3,849, cG50=4,388). The twenty-two autosomes are depicted as ideograms in the X-axis with the red band denoting the centromere position. The Y-axis for each row, represents the density of hypo-DMR regions present at each defined genomic window in the patient iPSCs and their corrected counterparts. Hypo-DMRs and CpG island (CGI) density (green) are partitioned into genomic windows of 1Mb and 1Kb, respectively.

To precisely determine the extent of DNA methylation recovery following CRISPR correction of *DNMT3B* mutations, we measured the number of hypo-DMRs in the corrected iPSC clones compared to WT1. We defined the ICF1 iPSC hypo-DMRs as restored when they were no longer differentially hypomethylated or if they were hypermethylated in the corrected clones. Out of the 26,031 and 22,727 hypo-DMRs in pR and pG iPSCs, 74.1% and 75% (n=19,291 and n=17,041) were rescued in both corrected iPSC clones of each patient iPSC (full rescue), respectively (**Table S3**), whereas 18.1% and 13.8% were corrected in only one iPSC clone (partial rescue) and 7.8% and 11.2% remained hypo-DMR in both corrected clones (no rescue). This again indicated that while the majority of the hypomethylated sites were amenable to correction by the restored DNMT3B, certain genomic regions resist rescue in all corrected clones.

### ICF1 DNMT3B variants fail to methylate CGI-rich chromosomal domains

We next asked whether the loci affected by DNMT3B dysfunction, and those that are rescued compared to those that resist rescue, are associated with specific genomic regions. To this end, we explored the distribution of hypo-DMRs across the autosomal chromosomes in the ICF1 iPSCs and their corrected counterparts (**Fig. 1B**). Strikingly, pR and pG iPSCs displayed a consistent overlap between the chromosomal distribution of hypo-DMRs. A pattern of distinct, sharp peaks with a higher density at the centromeric/pericentromeric and distal/telomeric regions were detected across almost all chromosomes. DNA methylation was restored in all four corrected clones with highly comparable patterns along all chromosomes (**Fig. 1B**). However, persisting hypo-DMRs were evident, and these occurred frequently at regions with the highest density of hypo-DMRs in ICF1 iPSCs, as apparent at many distal chromosomal regions. Conversely, centromeric and pericentromeric regions appeared more efficiently remethylated, as we previously reported [28], despite being characterized by marked peaks of hypo-DMRs in patient iPSCs.

At certain regions, we found that DNA methylation was not restored to the same extent in all corrected clones. While both corrected clones cG13 and cG50 showed relatively high levels of *de novo* methylation, cR7 and cR35 clones exhibited higher variability in the extent of DNA methylation rescue (**Fig. 1B**). The relative degree of remethylation in the various corrected clones was consistent at multiple genomic regions, supporting the notion that while all corrected clones regained DNMT3B catalytic activity, the individual clones vary slightly in the methylation efficiency of the restored enzyme.

An example of a potentially phenotype-related hypomethylation was observed at the pericentromeric region of chromosome 2, where two large blocks of HSATII flanking the immunoglobulin *IGK* gene cluster were evidently affected in both ICF1 iPSCs (**Fig. S2A**). Similarly, but at the distal region of chromosome 14q, the immunoglobulin heavy chain (*IGH*) loci were severely hypomethylated (**Fig. S2B**). The failure of the disease-causing *DNMT3B* variants to methylate these two gene clusters is relevant in the context of the ICF immune phenotype, in particular with relation to the process of VDJ rearrangements in early B cells [32]. A further confirmation that these genomic regions are primary targets of DNMT3B arises from methylation profiles of DNMT3B-depleted human embryonic stem cells (early and late 3BKO HUES64 hESCs and shRNA 3BKD H1 hESCs[33–35]) (**Fig. S2A-B)**, in which the increased size of hypomethylation sites is evident, compared to WT hESCs tracks. The position of genomic sites in DNMT3B deficient hESCs overlap those of hypoDMRs in ICF1 iPSCs at the large gene cluster domains (**Fig. S2A-B)**.

Overall, chromosomal regions with elevated hypo-DMR density correlated with higher gene and CpG Island (CGI) content (**Fig. 1B**), as clearly demonstrated at chromosomes 17, 19, 20 and 22. The chromosomal pattern of hypo-DMRs in ICF1 iPSCs indicates that several large gene clusters are poorly targeted by the mutated DNMT3B variants. In addition, we observed consistent hypomethylation at particularly long genes, as for instance the *PTPRN2* gene at the distal region of chromosome 7 (**Fig. S2C)**. These regions exhibited diffuse hypomethylation in ICF1 iPSC, and similar methylation defects were noticeably observed in the DNMT3B deficient hESCs.

To explore whether the hypomethylation is due to the inability of the *DNMT3B* variants to bind the target regions, we determined by ChIP-Seq the genome-wide DNMT3B binding profile in ICF1 iPSCs in comparison with that in WT1 iPSCs. Among the 19,706 DNMT3B enriched peaks detected in WT1 iPSCs, we observed significantly reduced binding (p-adj < 0.0001) in both pR and pG iPSCs (**Fig. S3A)**. The decrease in DNMT3B binding is less prominent in pG, despite the fact that these cells carry one *DNMT3B* null allele. However, we previously showed by western blot analysis that DNMT3B protein levels are comparable in both patient iPSCs [28], suggesting a compensatory mechanism for DNMT3B expression levels in pG. We next studied DNMT3B binding in the isogenic pR and pG iPSCs, cR7, cR35, cG13 and cG50, in which the *DNMT3B* mutations were corrected [28], and found that DNMT3B binding had recovered in approximately 75% of the regions affected in ICF1 iPSCs (**Fig. S3A;** see ChIP-Seq data analysis in “Supplementary Data”). Thus, while the majority of the genomic targets acquired proper DNMT3B binding following mutation correction, up to 25% of the affected regions failed to do so. This suggests that the recruitment of the corrected enzyme to these specific regions is impeded by molecular obstacles which constitute an epigenetic memory of the disease state. The profile of hypomethylated regions coincided with regions showing significant loss or reduced DNMT3B binding in both pR and pG iPSCs compared to WT1 (**Fig. S3B**), with the exception of centromeric and pericentromeric regions. This discrepancy results from intrinsic technical limitations of the ChIP-seq analysis which compromises peak detection at these specific repetitive sequences. Nevertheless, DNMT3B is known to specifically target repetitive regions [4], and we previously reported that DNMT3B binding was lost at pericentromeric satellite repeats in ICF1 iPSCs and fully restored following gene correction [28]. The majority of hypo-DMRs fall within a distance of 25kb from reduced DNMT3B binding regions (**Fig. S4A**), suggesting that DNMT3B may exert its catalytic activity on target regions through long-range interactions. This is evident, for example, at the genomic region upstream of the *TCF3* gene **(Fig. S4B),** where we observed a cluster of hypomethylated regions in both pR and pG iPSCs, flanking the DNMT3B binding sites. Noticeably, a domain of hypomethylated regions was also present in DNMT3B null hES [33–35], confirming that this region represents a hotspot of hypomethylation specifically associated with DNMT3B LOF. As expected, the recovery of DNMT3B binding in corrected clones reflected the distribution of the hypomethylated sites in ICF1 iPSCs (**Fig. S3B**).

### Promoter and intragenic methylation of genes with roles in immune response is impaired in ICF1 iPSCs

CGI-rich regions, highly correlated with gene coding regions, were the most highly enriched elements in hypo-DMRs across all chromosome arms. We therefore proceeded to characterize the specific gene components affected by DNA methylation loss in ICF1 iPSCs. We found that approximately half of the hypo-DMRs were localized in gene bodies, distributed along all exons, introns and 3’-UTRs, while about 15% mapped to gene promoters and 35% were present in intergenic regions (**Fig. 2A**). When we studied the correction of these gene-associated hypo-DMRs, we found that they are proportionally distributed along the various features (**Fig. 2A**).

**Figure 2.**
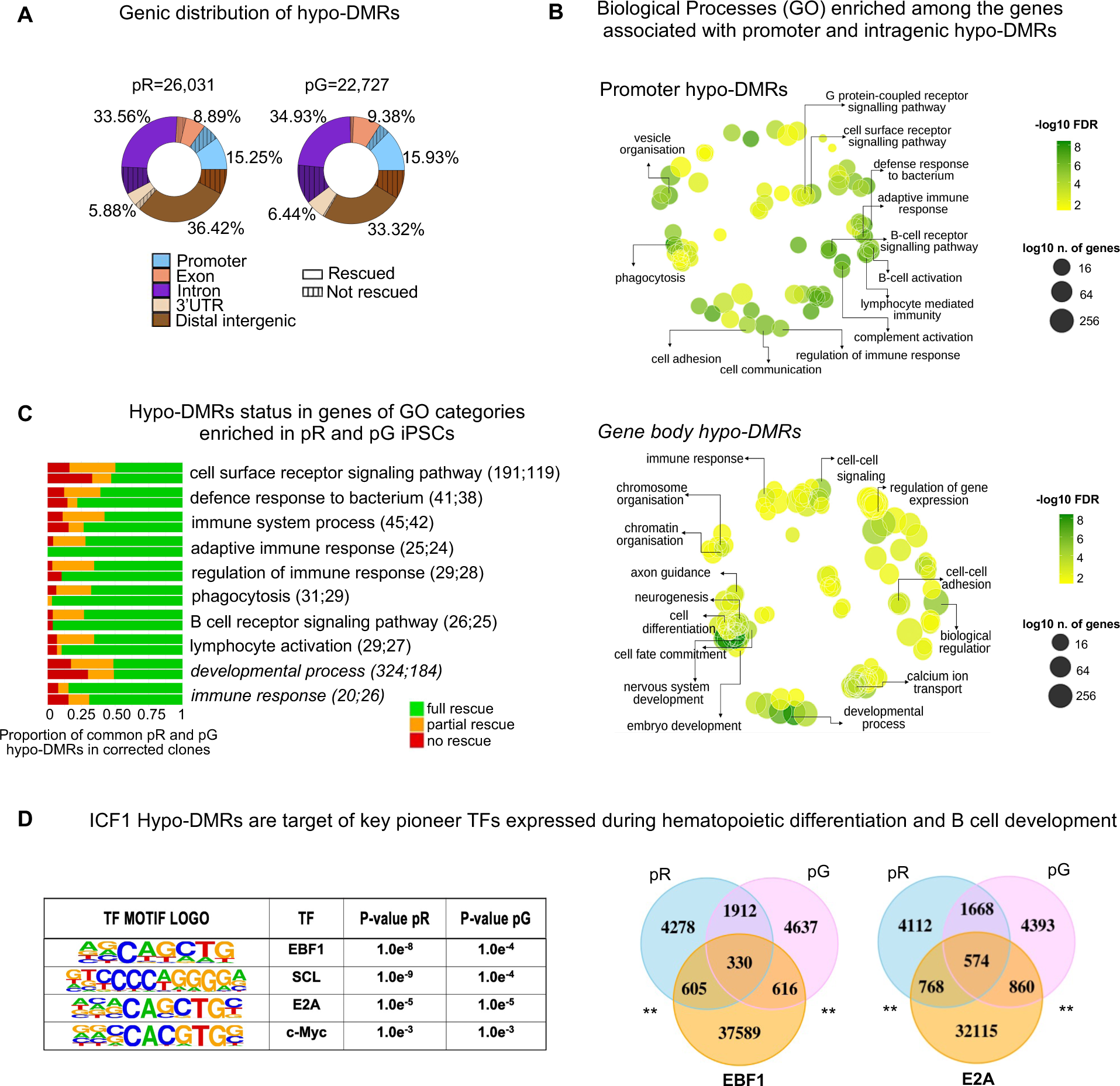
DNMT3B shapes CG methylation patterns at genes with functions in innate and adaptive immune responses. **(A)** Donut charts depicting the percentage distribution of hypo-DMRs in pR and pG, annotated to various genic regions. Striped areas: hypo-DMRs resistant to hypomethylation correction. **(B)** Gene Ontology terms (GO-BP) enriched for genes annotated to promoter and intragenic hypo-DMRs common to both ICF1 iPSCs (pR=1328 and pG=2336) are visualized as a Multidimensional Scaling scatter- plots. The color scale represents the -log10 of the adjusted p-value (Benjamini-Hochberg with False Discovery Ratio correction < 0.01; BH-FDR) of the GO-BP term. The dot size represents the log10 of the number of hypo-DMR associated genes in each GO-BP term. **(C)** Bar plot depicting the proportion of hypo-DMRs associated with genes enriched in each GO-BP category that are fully, partially or not rescued in corrected iPSCs clones. For each GO-BP, the upper and lower bars correspond to the hypo-DMRs status in pR and pG, respectively. GO-BP terms include genes with hypo-DMRs in promoters (roman) and in gene bodies (italic). The number of hypo-DMRs followed by the number of genes enriched in the GO-BP term is indicated within the brackets. **(D)** Left, Motif enrichment analysis of known Transcription Factors (TF) at hypo-DMRs with reduced DNMT3B binding (DERs). The output displays the predicted motif sequence and the p-value significance for ranking the motif enrichment at pR and pG hypo-DMRs. Right, Venn diagrams representing the overlap between ICF1 hypo-DMRs with enriched target regions of EBF1 and E2A TFs (peaks obtained from ENCODE ChIP-seq datasets of WT LCLs).

To investigate the functions of genes that display hypomethylation at promoters and intragenic regions in ICF1 iPSCs, we performed gene ontology (GO) analysis. Consistent with the immune phenotype present in ICF1 patients, we detected a significant enrichment of biological processes related to the regulation of the immune response. In particular, among genes with promoter hypomethylation we found regulators of both innate and adaptive immunity (**Fig. 2B; Table S3**). Biological processes related to cell fate commitment, embryonic development and neurogenesis were predominantly associated with genes hypomethylated at their bodies, consistent with the recognized role of intragenic CGI methylation in developmental regulatory genes [36, 37]. Remarkably, the majority of the immune response-related hypo-DMRs exhibited full recovery of normal DNA methylation patterns following correction of *DNMT3B* mutations (**Fig. 2C**).

The functional categories of the hypomethylated genes enriched in pR and pG iPSCs, resembled those of hypomethylated genes detected in lymphoblastoid cell lines (LCLs) derived from two unrelated ICF1 patients previously analyzed by our group using WGBS and RRBS [20]. We therefore exploited the methylation data of ICF1 iPSCs to evaluate whether the aberrant epigenetic signature generated during early developmental stages in ICF1 embryos persists in ICF1 patients’ somatic cells. Interestingly, when comparing the global methylation profiles of ICF1 iPSCs and LCLs [18, 20], we detected a significant overlap between hypo-DMRs in both cell types (pR and pG compared to WGBS- ICF1p1 and RRBS- ICF1p1 and ICF1p2; p-adjusted < 0.001) **(Fig. S5A)**. A significant similarity result was also observed when we analyzed the methylation data of primary blood samples from ICF1 patients [38]. Indeed, hypomethylated regions in ICF1 iPSCs overlapped, or were positioned within a distance of maximum 2kb (p-adjusted < 0.001), with those detected in ICF1 whole blood DNA **(Fig. S5B)** [38]. These results support the notion that a significant fraction of the abnormal methylation detected in ICF1 somatic cells arises during *de novo* establishment of DNA methylation around the implantation stage.

Highly relevant to the agammaglobulinemia characterizing ICF syndrome patients is the hypomethylation of the sequences encompassing the immunoglobulin heavy chain *IGHA1, IGHG1 and IGHG2* genes, which is similarly observed in ICF1p1 LCL WGBS (**Fig. S5C)**. Class-switch recombination (CSR) at the *IGH* locus is controlled by two complex regulatory regions (3’RRs) that are located downstream to the constant *IGHA1* and *IGHA2* genes. Noticeably, DNA methylation levels were reduced in the area encom these regulatory regions, particularly at the 3’RR nearby the *IGHA1* gene in both ICF1 iPSCs (**Fig. S5C**). In addition, conserved cis-element regions corresponding to switch regions (S) [39] were hypomethylated in ICF1 iPSCs. Following gene editing, the corrected DNMT3B was able to remethylate most of these regions (**Fig. S5C**), thereby supporting the prediction that gene correction might reverse epigenetic defects and consequential defects in the recombination process.

Additional immune response-related genes hypomethylated in ICF1 iPSCs and amenable to correction include tyrosine-protein kinases (*SRMS*), E3 ubiquitin-protein ligase TRIM proteins and Rho GTPase proteins *(RAC1)* (**Fig. S5D and Table S3**). Among the genes affected by promoter and/or intragenic hypomethylation in both patient iPSCs, we also detected key transcription factors and proteins regulating hematopoietic differentiation such as *RUNX1, TAL1, GATA2* and *CSF1R,* and early-stage regulators of B-lineage programming, such as *TCF3 (E2A), EBF1* and *EBF4* (**Fig. S5D and Table S3**). Moreover, genes involved in lymphocyte activation and the B-cell receptor signaling pathway *(SH2B2, VAV1, NFATC2, PLCG2, PRKCB)* were also hypomethylated. In addition to B-cell specific genes we detected hypomethylated genes with functions in T cell co-stimulation and in T cell receptor signaling pathways (*TNFSF14, CD28, AKT1, CARD11)* and the complement cascade (*C3*). Biological processes related to cell adhesion and migration (*TNXB, ITGA7, L1CAM*), as well as cell proliferation and differentiation were also enriched and include several genes (*FGF17, WNT5B*) (**Table S3**). Collectively, we identified genes that are early direct targets of DNMT3B-dependent *de novo* methyltransferase activity, whose hypomethylation might be functionally relevant in the context of the ICF immunodeficiency phenotype.

The hypomethylation patterns at the immune-related genes and their regulatory regions are frequently mirrored in ICF1 LCLs, strongly suggesting that these methylation profiles in somatic ICF1 cells are pre-marked early during embryonic development.

Additionally relevant to ICF1 abnormal phenotypes, motif enrichment analysis for transcription factor (TF) binding sites at ICF1 iPSCs hypo-DMRs highlighted binding motifs of pioneer TFs important for hematopoietic differentiation and B cell development, such as E2A and EBF1 [40], SCL/TAL1 [41], c-MYC [42] **(Fig. 2D and Table S3)**. Further evidence supporting the hypothesis that the hypo-DMRs might be target regions of these TFs emerged when we compared the hypo-DMRs with published TFs ChIP-seq datasets in wild-type hematopoietic stem and progenitor cells (HSPCs; [43]) and LCLs (ENCODE). We found a significant overlap between the hypo-DMRs in pR and pG ICF1 iPSCs and the corresponding TF peaks (**Fig. S5E**). Additionally, we identified by this analysis binding motifs for other TFs, such as ZIC2, ZIC3, LHX3, TCF12 and ASCL1, all which play central roles in neural differentiation and nervous system development [44–48], and several members of the DLX family of genes involved in craniofacial patterning and morphogenesis [49].

Collectively, these findings imply that abnormal hypomethylation at these TF targets might affect their recruitment, when expressed, during cell differentiation, thereby interfering with the normal regulation of tissue-specific gene expression programs during development of ICF1 embryos.

### Transcriptome profiling reveals minor changes in ICF1 iPSCs which correlate with an undisrupted pluripotent state

DNA methylation reduction in ICF1 iPSCs was frequently observed in proximity to gene regulatory elements, such as CGI-rich promoters and enhancers, changes that could potentially lead to disrupted gene expression during differentiation. Therefore, we further explored these classes of regulatory sequences in iPSCs carrying *DNMT3B* mutations. We intersected the hypo-DMRs in ICF1 iPSCs with 54,091 human regulatory elements reported in the GeneHancer (GH) database of promoters and enhancers and their inferred target genes [50]. Interestingly, hypo-DMRs in both pR and pG iPSCs significantly overlapped with, or were in close proximity to GH enhancers and promoters (6,063 and 6,214, respectively; p-adjust < 0.001), suggesting that DNMT3B LOF impairs the normal methylation of these regulatory sequences and their proximal flanking regions **(Fig. S5F,G**).

To investigate whether hypomethylation at these regions may directly affect gene expression in iPSCs, we performed RNA-Seq in WT1, ICF1 iPSCs and corrected clones and compared the datasets of gene expression and DNA methylation. The changes in global gene expression levels, as detected by RNA-Seq, were limited to 344 genes that were differentially expressed (DE; pp > 0.9) in both ICF1 iPSCs compared to WT1 iPSCs **(Fig. S6A and Table S4).** Among the deregulated genes in each ICF1 iPSCs, about 27% were either fully, partially or slightly rescued in their respective corrected clones **(Fig. S6B)**. Notably, out of the large group of genes hypomethylated at regulatory regions (CGIs, GH promoters and enhancers; pR=3178 and pG=3319), only a limited fraction of them demonstrated altered expression in ICF1 iPSCs (n=173 in pR and n=144 in pG, pp > 0.9), and we found no enrichment of these genes within the group of genes demonstrating the most severe hypomethylation (**Fig. S6C)**. Thus, DNA hypomethylation is insufficient by itself to disrupt global gene expression, and the effect of the epigenetic abnormalities present in stem cells is likely postponed to later developmental stages at which the affected genes are normally expressed. Consistently, several genes hypomethylated in both ICF1 iPSCs and LCLs, such as *LINC00221, IGHG1* and *RUNX1,* were differentially expressed only in ICF1 LCLs compared to WT LCLs **(Fig. S6D)**. Within the *IGH* gene cluster, in addition to *IGHG1,* the immunoglobulin genes *IGHA1, IGHG3* were hypomethylated and abnormally silenced in ICF1 LCLs, whereas the expression levels of *IGHM* gene was elevated [17, 18, 20, 51]. None of these genes were transcribed in ICF1 and WT iPSCs.

Nevertheless, it is important to note that among the hypomethylated genes whose expression was altered in ICF1 iPSCs, we could detect several neural and immune-related genes, which were similarly up- or down-regulated in ICF1 LCLs (for example *CD74, PLA2G4C)* (**Fig. S6E)** [18, 20, 51]. This suggests that the expression of a small subset of genes required for later immune and neural functions may be disrupted already at very early stages of development. In addition, when relaxing the threshold of the posterior probability for defining differential gene expression to > 0.8, we detected genes involved in B and T cell receptor signaling pathways *(VAV1, PRKCB)* as well as additional pathways contributing to the immune phenotype, such as signaling by Rho GTPases (*RAB17, RGS9*) and cytokine signaling in the immune system (*SQSTM1, TRIM14;* **Table S4**). Several deregulated genes were also functionally involved in neuron differentiation, nervous system development and axon guidance *(DPYSL5, RET* and *EFNB1).* Taken together, our results demonstrate that, although the gene expression status in ICF1 iPSCs is mostly undisrupted, it encompasses epigenetic and transcriptional alterations associated with ICF syndrome phenotypes.

Promoter hypomethylation is most commonly correlated with gene upregulation (**Fig. 3A and Table S4)**. We detected several genes associated with germline functions that were ectopically activated in both ICF1 compared to WT1 iPSCs. The overexpression of germline genes was described as a distinctive molecular characteristic of ICF1 patient blood when compared to ICF2-4 blood samples [38]. Some of these genes regained partial or full methylation in the corrected clones, as exemplified by the *RNF212* and *PTPN20* genes **(Fig. 3B,C)**. The CGI-associated hypo-DMR of the *RNF212* gene was rescued in the corrected clones of both ICF1 iPSCs, and consistently, its abnormal expression was reversed in all four clones. An alternative promoter serving as a TSS of an additional transcript isoform of *RNF212* similarly exhibited a hypo-DMR in ICF1 iPSCs, and was efficiently rescued in the corrected clones **(Fig. 3C and S7A)**. Strikingly, the recovery of proper gene expression levels is highly correlated with the degree of DNA methylation recovery, which in turn depends on the severity of hypomethylation in ICF1 iPSCs. One such example is the *TSPYL5* gene which exhibits high-score hypo-DMRs in ICF1 iPSCs. Persisting hypomethylation in the corrected clones of pG iPSCs is clearly correlated with remaining high expression levels, while partial correction of hypomethylation in the cR35 clone restored transcription to levels close to normal **(Fig. 3D and S7A)**. Taken together, these results indicate that preserving promoter CGI methylation is crucial for the repression of germline genes and, more importantly, that the recovery of normal DNA methylation patterns is sufficient to reestablish normal expression levels.

**Figure 3.**
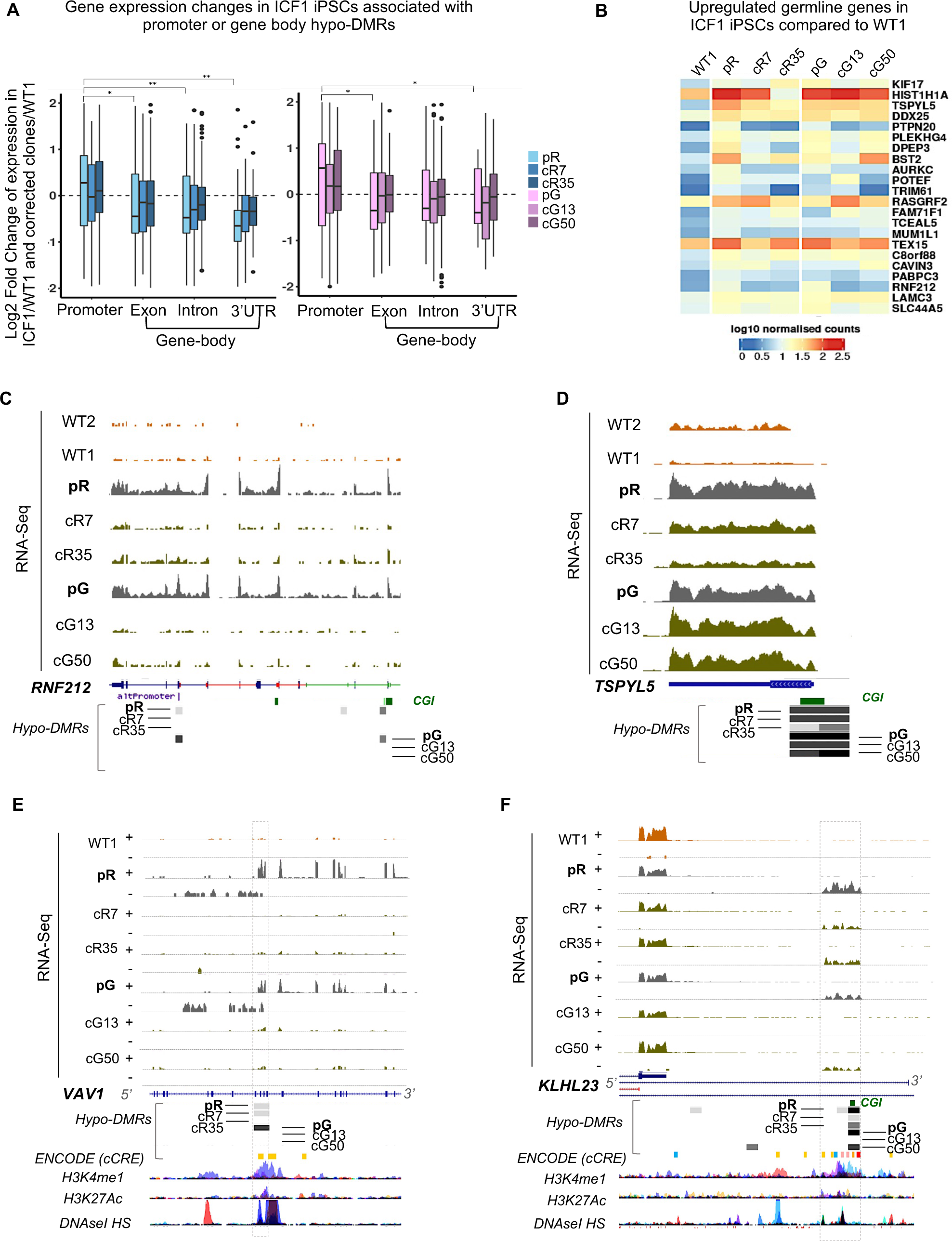
Transcriptional deregulation associated with promoter and intragenic hypo-DMRs in ICF1 compared to WT iPSCs. **(A)** Boxplot representation of the log2 fold change of the differentially expressed genes (pp > 0.8) with hypo-DMRs associated with regulatory elements (CGIs and GH elements) in pR (n= 6063) and pG (n= 6214) iPSCs and their respective corrected clones compared to WT1 iPSCs. The X-axis shows the gene feature category, and the Y-axis denotes the log2FC of the genes in each ICF1 iPSC vs WT1 iPSCs. The statistical significance of the differences in the log2FC of genes with hypo-DMRs annotated to promoters compared with genes containing hypo-DMRs annotated to other gene features was calculated using a non-parametric two samples two-sided Wilcoxon test with BH-FDR correction (*=FDR < 0.01 and **= FDR < 0.001). **(B)** Heatmap representing the expression levels of germline-specific genes hypomethylated and upregulated (pp > 0.8) in ICF1 iPSCs. The scale denotes the log10 of normalized counts of genes compared between ICF1 iPSCs and corrected iPSCs to WT1. **(C, D)** Genome browser views of the expression profiles of the hypomethylated germline-specific genes *RNF212* and *TSPYL5*. Grey boxes represent hypo-DMRs in ICF1 and corrected iPSCs clones compared to WT1 and WT2. The coverage tracks display the RNA-seq expression level of the genes in WT1 and WT2 internal and external controls, ICF1 and corrected iPSCs. **(E, F)** Genome browser views of *VAV1* and *KLHL23* gene regions. The strand-specific RNA-seq expression coverage tracks of WT1, ICF1 and corrected clones are shown. The gene and the intragenic enhancer-like region are distinguished by their expression in opposite strands. The hypo-DMR overlapping the enhancer-like region are shown below. Underneath, the yellow boxes denote ENCODE cCRE enhancers followed by the H3K4me1, H3K27Ac and DNAseI HS tracks.

In addition to upregulated gene expression associated with promoter hypomethylation, we observed deregulated genes with abnormal gene body DNA hypomethylation. In accordance with the previously described role of intragenic CG methylation in expression regulation [22, 25], we found that genes with reduced methylation in exons, introns and 3’UTRs were downregulated in pR and pG iPSCs (**Fig. 3A and Table S4)**. Remarkably, some downregulated genes regained normal intragenic DNA methylation upon correction of *DNMT3B* mutations, and concomitantly their transcription increased and approached normal levels (**Fig. 3A)**. For example, the downregulated *TMEM132D* gene exhibits multiple intragenic hypo-DMRs that appear less efficiently corrected in pR compared to pG. Accordingly, the transcriptional downregulation is more pronounced in pR than in pG and partial recovery of normal expression is evident only in cG13 and cG50 **(Fig. S7B)**. These findings support the view that gene body DNMT3B-dependent DNA methylation has a direct positive influence on transcript abundance in human iPSCs.

When further exploring the transcriptional changes in ICF1 iPSCs, we observed that several upregulated genes initiate from hypomethylated intragenic sequences, and these aberrant TSSs frequently overlap with enhancer-like regions from public databases. For example, the intragenic enhancer-like regions spanning exons 10-13 of the *VAV1* gene were targets of hypomethylation, and activated a bidirectional cryptic transcription initiation site on both strands in ICF1 iPSCs (**Fig. 3E)**. This abnormal transcription is reduced in all corrected iPSCs concomitantly with hypo-DMR rescue in all but clone cR7. However, when looking at the overall methylation track of the region, we observed partial correction of the methylation level in both cR7 and cR35 compared to pR (**Fig. S7C)**. At the *KLHL23* gene locus, hypomethylation and activation of an intronic enhancer-like region was associated with slight downregulation of the host gene in both ICF1 iPSCs (**Fig. 3F and S7D).** Full recovery of methylation occurred solely in the cG13 corrected clone, and this correlated with complete loss of the cryptic enhancer-like region transcription (**Fig. 3F**).

However, this change in cG13 had no direct effect on the recovery of normal *KLHL23* transcript levels, implying that the interplay between the upregulated enhancer-like regions and downregulated transcription of *KLHL23* involves complex regulatory mechanisms. These latter results demonstrate that DNMT3B is critical in preventing the activation of methylation-sensitive enhancer-like regions that can lead to cryptic intragenic initiation sites or perhaps influence the transcription of proximal genes in the proper developmental window.

### DNMT3B-variants fail to maintain normal DNA methylation level at boundaries of DNA methylation valleys and CGIs

The large hypomethylated domains in ICF1 iPSCs, motivated us to further explore whether DNMT3B LOF affects the status of DNA methylation valleys (DMVs) identified in ES cells [52]. DMVs are defined as genomic regions devoid of DNA methylation, highly conserved among different cell lines and species.

Interestingly, we found that out of 639 DMVs, 223 and 211 DMVs overlap or are in close proximity (+/- 5Kb) with the hypo-DMRs in pR and pG iPSCs, and 139 are shared between both patient iPSCs (p-adjust < 0.001). Strikingly, the normally methylated DMV borders were frequently hypomethylated in ICF1 iPSCs and in DNMT3B deficient hESCs, leading to expansion of the unmethylated valleys in both directions, as shown at the *ICAM5* gene in **Fig. 4A,B.** This indicates that DNMT3B contributes to preserving the methylation state of DMVs borders in normal pluripotent stem cells.

**Figure 4.**
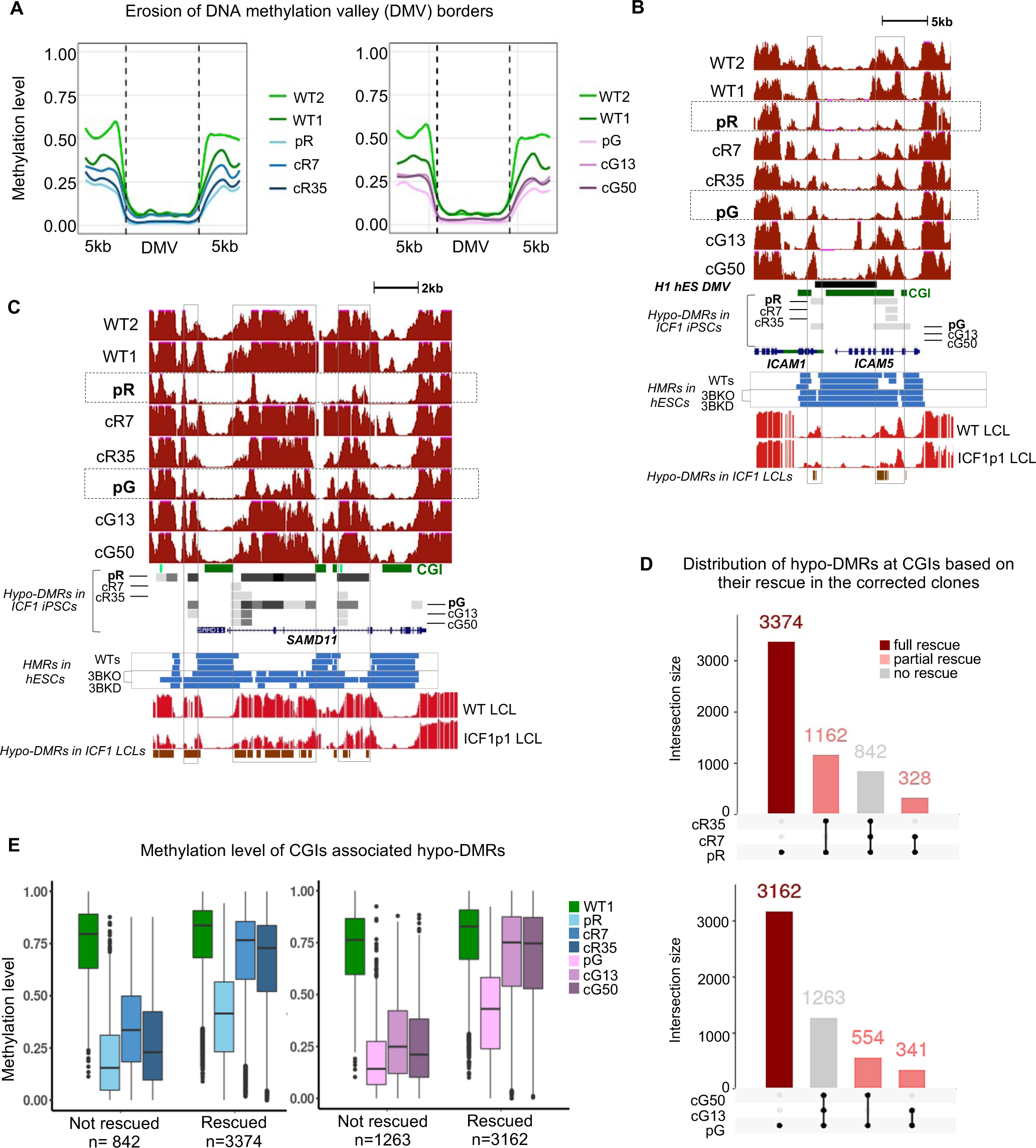
DNMT3B fails to maintain methylation levels at boundaries of DMVs and CGIs in ICF1 iPSCs. **(A)** Plots of average weighted CG methylation levels across DNA methylation valleys (DMVs) and flanking 5kb regions in ICF1 and corrected iPSCs clones in comparison to the WT1 and WT2 iPSCs. The 0 to 1 scale at the Y-axis denotes the CG methylation level (expressed as ratio of the number of Cs over the total number of Cs and Ts) measured in each bin. **(B)** A genome browser view of a DMV in the *ICAM5-ICAM1* gene loci, representing an example of erosion of valley boundaries in ICF1 iPSCs and LCLs compared to the WT counterparts. The black box represents the DMV, as identified in H1 hESC [52] and grey boxes indicate the hypo-DMRs in ICF1 and corrected iPSCs. The underneath blue tracks illustrate the hypomethylated regions (HMRs) in WT human embryonic stem cells (hESCs) from H1, HUES9 and H9 lines, followed by DNMT3B-KO hESCs (early and late passage DNMT3B-KO; 3BKO) and shRNA DNMT3B-KD hESCs; 3BKD). Bright red tracks denote methylation coverage measured by WGBS in ICF1p1 LCLs and WT LCLs and brown boxes underneath represent hypo-DMRs in ICF1p1 LCLs. **(C)** A genome browser view of CGIs in the *SAMD11* gene, representing an example of methylation loss at CGI edges in ICF1 iPSCs and LCLs compared to WT counterparts. The tracks are shown as in B. **(D)** Upset plots showing the hypo-DMRs at CGIs (+/-2kb) in pR (top) and pG (bottom) iPSCs and their corrected counterparts. Vertical bars represent the intersection size between the hypo-DMRs present in each iPSC sample. Black dots in the three rows below each panel represent the hypo-DMRs present in each intersection. Full, partial and no rescue of ICF1 hypo-DMRs indicate remethylation in both, single, or none of respective corrected clones. **(E)** Boxplots representing the distribution of methylation levels at CGI-associated hypo-DMRs in ICF1 iPSCs which remain hypomethylated (not rescued) and those that are rescued in the corresponding isogenic clones.

To further investigate whether the erosion of DMV borders may generally affect promoter and intragenic CGIs, we plotted the average methylation profiles of CGIs at the tertile with the lowest methylation level in WT1 iPSCs. We observed hypomethylation at CGI-flanking regions (+/-5kb), suggesting that DNMT3B LOF results in the expansion of hypomethylation at many unmethylated CGIs at both edges, and not only at the subset included in DMVs **(Fig. S8A)**. To further explore this observation, we intersected the hypo-DMRs detected in ICF1 iPSCs with the list of UCSC CGIs and detected approximately 2,600 hypomethylated regions overlapping with CGIs out of 26,031 and 22,727 hypo-DMRs in pR and pG iPSCs, respectively **(Fig. S8B)**. However, interestingly, when CGIs were extended by +/-2kb, their intersections with hypo-DMRs doubled in both pR and pG iPSCs (p-adjust < 0.001), indicating that the flanking regions of CGIs constitute major hypomethylated elements in ICF1 iPSCs.

The erosion of edges of unmethylated CGI regions in pR and pG iPSCs is particularly evident in a subgroup of CGIs, in which the nearby hypomethylated CGIs merge into a larger hypomethylated domain, demonstrated for example at the *SAMD11* gene locus **(Fig. 4C).** This pattern of hypomethylation was clearly observed also in ICF1 LCLs, indicating that this abnormality is disease-specific and not associated with specific *DNMT3B* variants (**Fig. 4C)**. Also, it is evident in DNMT3B deficient hESCs, confirming that it is a direct target of DNMT3B during *de novo* methylation. Notably, the majority of the extended hypomethylated region present between the *SAMD11* CGIs was remethylated to varying degrees in all patient corrected iPSC clones. However, interestingly, the edges of these regions exhibited higher resistance to *de novo* methylation (**Fig. 4A,C)**.

We next wanted to measure the overall level of methylation recovery at hypo-DMRs localized to CGIs. We observed that approximately 60% of this subgroup (3374 and 3162 in pR and pG, respectively) were no longer hypomethylated in both corrected clones of pR and pG iPSCs (**Fig. 4D)**. Remarkably, we identified 2,485 hypo-DMRs at CGIs that were commonly hypomethylated in both patients’ iPSCs and 1,161 of them regained a normal methylation profile in all corrected counterparts. On the other hand, a subgroup of CGI associated hypo-DMRs was resistant to remethylation in all four corrected clones (18%). We then searched for molecular characteristics shared by the persistently hypomethylated regions that may explain their failure to regain methylation by the corrected DNMT3B. While the size, density and position of CGIs (promoter or intragenic) do not appear to influence the ability to regain methylation in corrected iPSCs, we found that the resistant regions were all characterized by the lowest methylation levels identified in patient iPSCs compared to WT1 (**Fig. 4E**). Collectively, these results suggest that the mutant-DNMT3B in ICF1 iPSCs fails to maintain proper methylation level at CGI boundaries, and that the corrected DNMT3B can fully restore the normal methylation signature only at CGI sites that display milder hypomethylation in patient iPSCs.

We then evaluated the percentage of hypomethylated GH promoters and enhancers that reacquire proper DNA methylation following *DNMT3B* correction and found that similar to the observed at CGIs, they amounted to 68% and 67% in both corrected clones of pR iPSC and pG iPSCs, respectively (**Fig. S8C**). The *WDR97* gene represents an example of methylation loss at enhancer regions (ENCODE cCRE and GH) in pR and pG iPSCs, and ICF1 LCLs (**Fig. S8D**). Again, we found that the original level of methylation of these regions in patient iPSCs influenced the degree of methylation recovery (**Fig. S8E**).

These results confirm that at gene regulatory regions as well, the loss of DNA methylation beyond a certain threshold, impedes the salvage of normal methylation patterns, despite the recovery of normal DNMT3B enzymatic activity.

### Corrected DNMT3B cannot overcome the highest abnormal increase of H3K4me3 at the fraction of persisting hypo-DMRs

Following CRISPR editing, we observed partial recovery of DNA methylation in all corrected clones of both patient iPSCs, with the majority of genomic regions regaining WT methylation levels. This confirms that the corrected DNMT3B protein restored its enzymatic activity, and suggests that the feature common to regions resistant to *de novo* methylation is their inability to recruit the rescued DNMT3B. We previously showed that subtelomeric regions with persisting hypomethylation in the corrected ICF1 iPSCs were characterized by high levels of H3K4me3 as result of residual epigenetic memory, which inhibited the regain of normal DNA methylation [28].

To assess whether similar to subtelomeric regions, high H3K4me3 levels interfere with DNA methylation rescue at non-repetitive regions throughout the genome, we examined the epigenetic landscape of hypo-DMRs, both rescued and unrescued, which are associated with the CGIs or GeneHancer (GH) regulatory elements previously examined (**Fig. 4E and S8E**). These subgroups of hypo-DMRs were merged after removing duplicates, and analyzed for enrichment profiles of H3K4me3, H3K36me3 histone marks and DNMT3B binding obtained by ChIP-Seq in WT1, ICF1 iPSCs and their corrected counterparts **(Fig. 5A and S9A).** The analyzed hypo-DMRs were divided in the complex heatmap into four groups based on their methylation level in ICF1 iPSCs.

**Figure 5.**
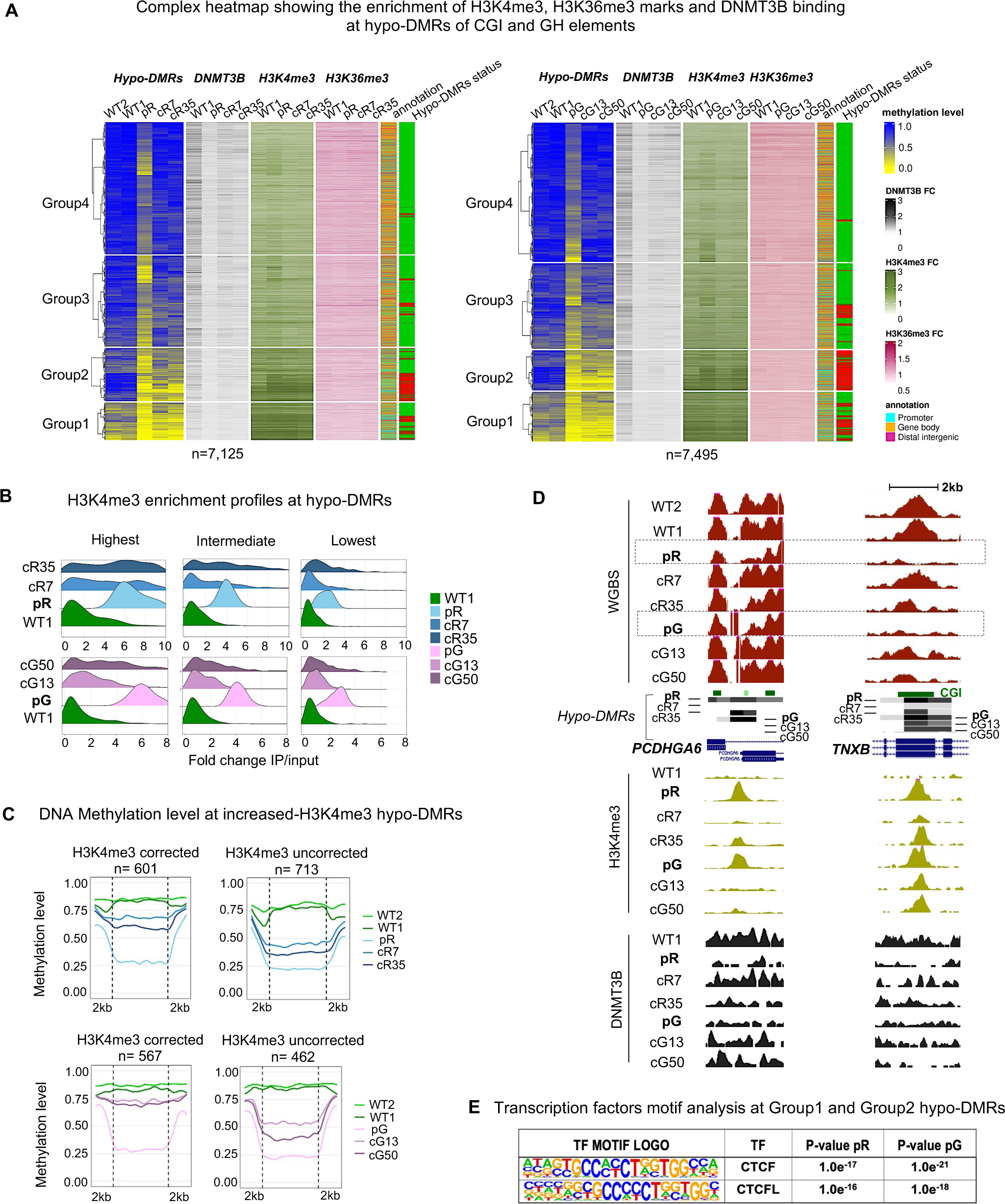
Hypo-DMRs persist at regions with the highest aberrant increase of H3K4me3 which prevents DNMT3B binding. **(A)** A complex heatmap showing the integration of ChIP-Seq data of DNMT3B binding, H3K4me3 and H3K36me3 marks with methylation levels of hypo-DMRs at CGI and GH elements. The gene annotation of the hypo-DMRs and their status of correction (green= rescued, red= not rescued) are depicted on the right. The hypo-DMRs are clustered using k-means into four groups based on their mean methylation levels in ICF1 iPSCs. FC=Fold change (IP/input) of the ChIP-seq enrichment calculated at the hypo-DMRs. **(B)** Ridgeline plots depicting the distribution of the FC of H3K4me3 enrichment at increased H3K4me3 DERs (Differentially Enriched Regions) proximal to ICF1 hypo-DMRs. The increased H3K4me3 DERs in pR and pG were divided into three groups based on the FC ranking levels in ICF1 iPSCs, and the FC was calculated for the respective corrected clones at the DERs. **(C)** Plots of average CG methylation levels across hypo-DMRs associated with increased H3K4me3 DERs in ICF1 iPSCs. The left panels include DERs with increased H3K4me3 in which the H3K4me3 levels are restored to normal levels in both corrected clones of pR (top) and pG (bottom), while the right panels include increased H3K4me3 DERs which persist in corrected clones of pR and pG. **(D)** Genome browser views of representative regions characterized by abnormal increase of H3K4me3 level at hypo-DMRs in ICF1 iPSCs. Following DNMT3B corrections, the H3K4me3 levels are fully or partially reverted in corrected clones together with hypomethylation at the *PCDHGA6* gene, while it persists at the *TNXB* gene. Green and black tracks represent H3K4me3 and DNMT3B binding enrichment levels. **(E)** Motif enrichment analysis of known Transcription Factors (TF) at hypo-DMRs with reduced DNMT3B binding (DERs) and increased H3K4me3 levels. The panel corresponds to motifs enriched at the subset of hypo-DMRs corresponding to Group 1 and Group 2 in pR and pG iPSCs, as shown in Figure 1B.

The hypo-DMRs of Group1 and Group2, demonstrated the lowest levels of DNA methylation level in ICF1 iPSCs and a limited recovery of DNMT3B binding in the corrected clones, while the hypo-DMRs in Groups 3 and 4, demonstrated recovery of DNMT3B binding in the corrected clones and almost full rescue of DNA methylation levels (**Fig. 5A)**. Group1 and Group 2 hypo-DMRs were mostly annotated to promoters and showed the highest enrichment of H3K4me3 in all four iPSCs compared to Group 3 and 4 hypo-DMRs that were mostly annotated to gene bodies. With the exception of Group1 hypo-DMRs, which already showed the initial highest H3K4me3 level, in all the other groups we clearly observed that regions with the lowest methylation level in ICF1 iPSCs correlated with the highest H3K4me3 increase, thereby reinforcing our previous findings at subtelomeric regions [28].

We additionally observed that while hypomethylation in pR and in pG iPSCs correlated with increased levels of H3K4me3, the level of H3K36me3 mark at hypo-DMRs was less affected in ICF1 iPSCs compared to WT iPSCs (**Fig. 5A**). This suggests that DNMT3B LOF does not strongly affect the activity of SETD2 histone H3K36 methyltransferase in ICF1 iPSCs and is consistent with the observations that deposition of H3K36me3 by SETD2 is an event upstream of DNMT3B mediated methylation [22, 23, 35, 53].

Noticeably, many regions abnormally enriched with H3K4me3 in ICF1 iPSCs did restore normal levels of this histone mark following correction of *DNMT3B* mutations **(Fig. 5A).** This is commonly observed in all corrected ICF1 iPSCs, particularly in regions belonging to Groups 3 and 4. These regions overall overlap with those showing rescue of normal DNA methylation levels (green in last column of the heatmap) and recovery of DNMT3B binding in the corrected clones. This correlation is also exemplified in corrected clones cR7 and cG13, which restored normal H3K4me3 levels more efficiently than cR35 and cG50, and in addition display higher degrees of DNA methylation and DNMT3B binding rescue.

To quantify the association between hypomethylation and increased H3K4me3 enrichment in ICF1 iPSCs, we intersected the 26,031 and 22,727 hypo-DMRs in pR iPSCs and pG iPSCs, respectively, with genomic regions exhibiting a significant increase in H3K4me3 level (differentially enriched regions=DERs) in pR (n=2,823) and pG (n=1,178) ICF1 iPSCs. We detected 1,442 and 1,081 hypo-DMRs overlapping, or in close proximity (+/-2kb), to increased H3K4me3 DERs in pR and pG, respectively (**Fig. S9B). However, all the significantly increased H3K4me3 DERs (2,823 and 1,178) were characterized by very low DNA methylation levels (**≤**0.25) in patient iPSCs(Fig. S9C,D**), indicating that H3K4 trimethylation mark significantly increases at regions where DNA methylation is reduced below a certain threshold.

To further explore H3K4me3 levels in WT, ICF1 and corrected clones at hypo-DMRs overlapping with increased H3K4me3 DERs, we divided them in three groups based on the H3K4me3 enrichment level in ICF1 iPSCs (**Fig. 5B**). Regions demonstrating the highest increase in H3K4me3 levels in pR and pG iPSCs were restored less efficiently to WT H3K4me3 levels in all corrected clones, compared to those showing an intermediate or low H3K4me3 increase in ICF1 iPSCs. Therefore, the subset of hypo-DMRs with the highest H3K4me3 increase in ICF1 iPSC are resistant to changes in this histone modification following DNMT3B correction. Out of 1,442 hypo-DMRs in pR and 1,081 in pG, this subset of regions included 713 and 462 hypo-DMRs, while 601 and 567 hypo-DMRs restored normal H3K4me3 levels in the corrected clones. Remarkably, the highly enriched H3K4me3 class of hypo-DMRs remained hypomethylated, while hypo-DMRs which restored normal H3K4me3 levels regained DNA methylation levels in the corrected clones to a much larger extent **(Fig. 5C).**

One region that exemplifies this situation encompasses the *PCDHGA6* gene from the protocadherin gene cluster that contains hypo-DMRs with abnormally high H3K4me3 levels in ICF1 iPSCs. In cR7, cG13 and cG50 corrected clones H3K4me3 levels were reverted back to normal, and simultaneously, the corrected DNMT3B was recruited and normal DNA methylation levels were restored. In the cR35 clone, where the H3K4me3 barrier was not entirely removed, the DNA hypomethylation was not corrected. On the other hand, at the *TNXB* gene where H3K4me3 levels are abnormally high and persisting in all iPSCs, DNMT3B binding and DNA methylation levels were not restored (**Fig. 5D and S9E)**.

The above results reveal that a subset of genomic regions can overcome the aberrant H3K4me3 increase and hence recruit the corrected DNMT3B protein to reestablish the normal DNA methylation pattern, while others are impervious to such a correction. We hypothesized that DNMT3B binding may be impeded at the resistant regions by additional bound factors with affinity to high H3K4me3 levels. To explore this possibility, we used the HOMER suite to perform motif enrichment analysis for transcription factor (TF) binding sites at all hypo-DMRs showing persistent loss of DNMT3B binding in combination with the highest H3K4me3 and lowest DNA methylation levels (belonging to Groups 1 and 2 shown in Figure 1B)(**Fig. 5E)**. We additionally analyzed hypo-DMRs from Groups 3 and 4 showing recovery of normal DNA methylation levels and DNMT3B binding. Noticeably, we observed a significant enrichment of binding motifs for CTCF and its paralogous BORIS (CTCFL) factors at high frequency within hypo- DMRs of Groups 1 and 2 (p < 1e^-16^) (**Fig. 5E and Table S5)**. Hypomethylated regions from Groups 3 and 4 showed a significant but lower enrichment of CTCF binding sites (p < 1e^-4^) with a lower frequency (6%) compared to Groups 1 and 2 (17%) (**Table S5)**. This analysis suggests that the failure to restore DNA methylation at these target regions might be promoted by the abnormal binding of the CTCF chromatin insulator in ICF1 iPSCs, that in turn might impede the recruitment of the corrected DNMT3B protein.

## DISCUSSION

The initial whole genome DNA methylation patterns are established around implantation by the action of the *de novo* DNMTs, with DNMT3B playing a major role compared to DNMT3A [3]. Later in development, during differentiation to specific lineages, tissue-specific DNA methylation patterns are formed. It is unclear to what degree disruptions in DNA methylation patterns during implantation in ICF1 embryos persist throughout development, and whether they can affect the pattern of methylation in differentiated cells. ICF1 patient-derived iPSCs mimic the initial developmental stage at which the mutated DNMT3B fails to properly methylate its genomic targets. Therefore, these cells are a valuable platform to identify genomic regions that are affected early in development, and to follow-up their behavior in ICF1 somatic cells.

CRISPR/Cas9 based correction of the pathogenic DNMT3B variants in ICF1 iPSCs enables to evaluate of the capacity to restore normal methylation levels. Previously, we demonstrated that following correction of two ICF1 iPSCs, normal methylation levels were fully restored at pericentromeric satellites, whose hypomethylation is a hallmark of ICF syndrome [13]. In contrast, abnormal DNA hypomethylation persisted at several subtelomeric regions, indicating that certain genomic regions resist *de novo* methylation by the corrected DNMT3B. Here we expanded our study to the non-repetitive fraction of the genome to characterize the abnormal epigenetic landscape of the two ICF1 iPSCs and their isogenic corrected clones, aiming to understand the molecular consequences of DNMT3B LOF throughout the entire genome and the feasibility of rescuing an abnormal epigenome arising in a human epigenetic disease. This is, to our knowledge, the first attempt to evaluate whether the sole correction of an enzymatic deficiency responsible for the molecular pathology of an inherited epigenetic disease, suffices to rescue genome-wide epigenetic perturbations.

Our comprehensive analysis detected highly comparable patterns of DNA hypomethylation in both ICF1 iPSCs, highlighting the preferential targets of DNMT3B across the genome. At the chromosomal level, we observed that in addition to the pericentromeric regions, hypo-DMRs were detected at distal regions of several chromosomes. Large domains of CGI-rich regions and gene clusters were also characterized by loss of DNA methylation (Fig. 1B). These abnormally low levels of DNA methylation were accompanied by a marked loss of DNMT3B binding (Fig. S3A), and overall, the chromosomal hypomethylation pattern clearly coincided with that of lost or reduced DNMT3B binding in both ICF1 iPSCs (Fig. S3B). Defective DNMT3B binding occurred despite the fact that the causative mutations are present in the C-terminal region encoding the catalytic domain, while DNA binding by DNMT3B is facilitated by a N-terminal domain. This confirms the biochemical evidence that DNA binding and enzymatic properties of DNMT3B during *de novo* methylation are tightly interconnected, and that mutations in the catalytic domain may perturb DNMT3B function at multiple levels [54–56]. Comparing the methylation defects in ICF1 iPSCs with that of DNMT3B knockout in hESCs provided further confirmation that the regions identified represent primary and direct DNMT3B targets methylated during the *de novo* wave of methylation.

A closer look at specific DNMT3B target regions revealed that large hypomethylated domains were frequently flanked by peaks of enrichment (Fig. S4B), suggesting that DNMT3B may exert its catalytic activity at target regions through long-range interactions. Three-dimensional chromatin structure and DNA methylation are implicated in multiple developmental processes [57], but the precise role of DNMT3B in these processes is not fully characterized. Recent observations suggest that in stem and progenitor cells long loops connecting even dozens of mega bases apart may be anchored in large DNA methylation “grand canyons” [58]. These regions, known also as DNA methylation valleys (DMVs), are defined as genomic regions devoid of DNA methylation, which are highly conserved among different cell lines and species [52]. Deletion of Dnmt3a in mouse hematopoietic stem cells led to the erosion of DNA methylation at canyon borders [59]. Interestingly, DNMT3B LOF in ICF1 iPSCs similarly affects methylation levels at DMVs edges [52], thus leading to DMV expansion. This is observed also at a significant number of CGIs and unmethylated enhancers throughout the genome. Therefore, DNMT3B contributes to preserving the methylation status of DMVs, CGIs and enhancer borders in WT pluripotent stem cells.

In addition to the abnormal methylation at heterochromatic regions, ICF1 is also characterized by gene expression deregulation due to DNA methylation changes [18–20, 51]. When we studied gene regions in ICF1 iPSCs we frequently observed abnormally low DNA methylation levels in proximity to gene regulatory elements, such as CGI-rich promoters, as well as at enhancer-like regions (Figs. 2-4).

Consistently, we observed a significant overlap between the hypo-DMRs identified in ICF1 iPSCs and those described in ICF1 patient peripheral blood and derived LCLs [20, 38]. However, we detected only mild transcriptional changes in ICF1 iPSCs, and the stability of the pluripotent state of the ICF1 iPSCs was consistently undisrupted. This suggests that preserving the proper methylation marks at gene regulatory elements in stem cells is essential for ensuring correct gene expression patterns at the relevant developmental windows. Thus, the major transcriptional consequences of aberrant hypomethylation of most gene regions are presumably postponed to later developmental stages in somatic cells at which the affected genes are normally activated or at an aberrant temporal window in the case of precocious gene activation.

When scrutinizing the genes specifically hypomethylated in ICF1 iPSCs, we observed a striking enrichment of genes involved in transcriptionally altered pathways in ICF1 LCLs, such as B cell receptor signaling, T cells co-stimulation, cytokine signaling in the immune system and receptor tyrosine kinase signaling [20]. This further supports the notion that abnormally hypomethylated genes displaying normal expression levels during implantation may manifest an abnormal transcriptional effect only at later developmental stages when additional required factors are present. A highly relevant example is the immunoglobulin *IGH* gene cluster, which is hypomethylated in ICF1 iPSCs and in ICF1 LCLs, but transcriptionally affected only in ICF1 LCLs and peripheral blood. Within the *IGH* cluster, the immunoglobulin *IGHA1*, *IGHG1*, *IGHG3* genes are aberrantly silenced in ICF1 LCLs, whereas the expression of *IGHM* and *IGHD* genes is variably affected [17, 18, 20, 51]. These transcriptional defects are strongly associated with the hypogammaglobulinemia caused by the defective terminal B-cell differentiation, and are the major cause of immunodeficiency in ICF syndrome patients [6, 16].

More recently, the loss of ZBTB24 and Lsh/HELLS functions, responsible for ICF2 and ICF4 subtypes, respectively, has also been associated with specific defects in immunoglobulin class-switch recombination and consequently immunoglobulin production and isotype balance [60, 61]. However, the exact molecular link between hypomethylation at the *IGH* cluster and *IGHG* and *IGHA* transcriptional repression in ICF1-4 patients is still unknown. Interestingly, recent studies in mice demonstrate that methylation patterns at the majority of immunoglobulin heavy chain locus cis-elements, including promoters, enhancers and insulators, are established early in development and faithfully maintained independently of B cell activation or transcription initiation at the constant genes [62]. Considerable hypomethylation in ICF1 iPSCs compared to WT was similarly detected across the *IGK* immunoglobulin kappa locus at the pericentromeric region of chromosome 2, in particular at two large blocks of HSATII repeats sequences flanking the group of exons encoding for *IGKVD* segments (Fig. S2A). Whether maintaining the proper methylation pattern of these satellite repeats is important during the immunoglobulin gene rearrangements has not been explored and requires further investigation.

Nevertheless, studies in mice demonstrated that Dnmt3a and Dnmt3b are critical for regulating the onset of Igκ light chain rearrangement during early B-cell development, and their deficiency causes precocious recombination of the immunoglobulin κ light chain [32].

Acquisition of hematopoietic and lymphopoietic cell identity requires strict epigenetic control to ensure stepwise activation of lineage-specific gene expression programs in multipotent progenitors to mature cells. In particular, an increasing body of evidence suggests that gene expression changes that direct hematopoietic and B cell differentiation are driven by both the action of transcription factors (TFs) as well as by DNA methylation changes. It has been suggested that several TFs that play key roles in hematopoietic and B cell fate decisions, such as E2A, EBF1 [40], SCL/TAL1 [41], and C-MYC [42], act as pioneer factors by binding DNA methylated regions and inducing local chromatin accessibility and demethylation of naive chromatin [63–67]. This crosstalk between TFs and the epigenetic machinery is required for establishing lineage-specific identity. Therefore, disruption of the normal methylation in ICF1 iPSCs at the target binding sites of these TFs might affect their activity during cell differentiation, thereby interfering with the normal regulation of lineage-specific gene expression programs. Interestingly, we found that a significant fraction of hypo-DMRs in ICF1 iPSC coincides with the target regions of these TFs in hematopoietic stem and progenitor cells and LCLs. However, further studies in ICF1 iPSCs differentiated towards hematopoietic progenitors are required to fully investigate this aspect.

Although regulation of gene expression is not globally disrupted in ICF1 iPSCs, we detected upregulation of several lineage-commitment genes, indicating that DNMT3B-mediated DNA methylation may prevent their ectopic expression. Moreover, we found that hypomethylation of intragenic methylation-sensitive enhancer-like regions can activate sites of cryptic transcription initiation, which frequently are corrected when the activity of DNMT3B is restored. The evidence that DNMT3B-mediated DNA methylation suppresses illegitimate transcripts from cryptic TSS in gene bodies was reported in several mouse and human ES cells, and recently assessed in Dnmt3b-deficient mice embryos, indicating that this function of DNMT3B is well conserved across species [3, 24].

The recent development of genome editing tools promotes new strategies for correcting genetic diseases. While there are genetic diseases in which the repair of an enzyme is likely to rescue the disease phenotype, it is unclear whether this is achievable in the case of an epigenetic disease in which the enzymatic disorder affects the epigenetic landscape throughout the genome, and thus the expression of multiple genes. It is also unclear whether the abnormal epigenome that forms as a consequence of DNA hypomethylation, as in ICF syndrome, is reversible. As iPSCs model the earliest developmental stage at which DNA methylation is disrupted, we took advantage of this platform to discover whether the rescue of the molecular abnormalities in ICF1 syndrome is feasible. Remarkably, upon CRISPR/Cas9 correction of the *DNMT3B* disease-causing variants, the majority (75%) of the genomic targets regained DNMT3B binding, and respectively, the majority of the hypomethylated sites in ICF1 iPSCs fully or partially restored WT iPSC methylation patterns. For example, the corrected DNMT3B restores most of the normal methylation pattern at the *IGH* locus (Fig. S2B). Additionally, several immune-related genes with functions in the innate and adaptive immune responses regained normal DNA methylation levels in the corrected clones (Fig. 2C), thereby indicating that these genes are pre-marked by DNMT3B activity at a very early stage of development.

Nevertheless, up to 10% of the affected regions failed to regain DNA methylation in both corrected clones, and 15% were corrected in only one iPSC clone, suggesting that the recruitment of the corrected enzyme to these specific regions is impeded by the epigenetic memory associated with the disease state in the parental cells.

Our findings suggest that binding of the corrected DNMT3B is inhibited at regions characterized by the most severe hypomethylation in patient iPSCs (Fig. 5A). These regions in most cases, also display the highest abnormal increase of H3K4me3. These findings are in agreement with our previous observations that subtelomeres resistant to *de novo* methylation following *DNMT3B* correction are those that manifest the highest degrees of DNA hypomethylation and H3K4me3 enrichment [28]. Interestingly, at regions in which H3K4me3 was elevated by a milder degree, the corrected DNMT3B protein could overcome the abnormal H3K4me3 levels and restore normal DNA methylation and H3K4me3 levels (Fig. 5B-D). Thus, there appears to be a certain threshold of H3K4me3 enrichment beyond which DNMT3B fails to bind its target regions and reestablish normal chromatin characteristics (Fig.6).

**Figure 6.**
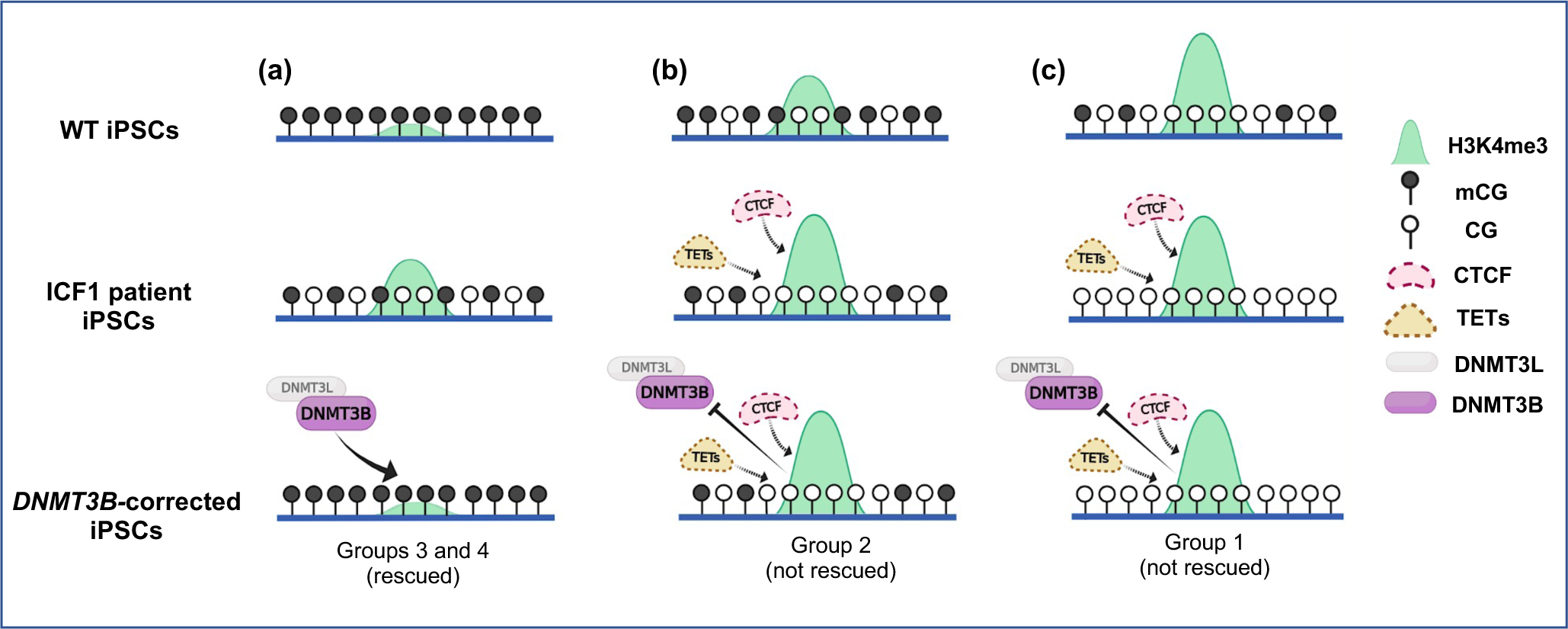
Model representing how abnormal H3K4 trimethylation influences the restoration of DNA methylation in corrected iPSCs. Graphical representation of hypo-DMRs identified in ICF1 iPSCs and clustered in Groups 1-4 based on their DNA hypomethylation level, as reported in Fig. 5A: **(a)** rescued hypo-DMR of Groups 3 and 4, showing high methylation and low H3K4me3 levels in WT iPSCs, mild hypomethylation and H3K4me3 increase in ICF1 iPSCs, and regain of normal DNA methylation and H3K4me3 levels upon restoration of DNMT3B activity by gene editing; **(b)** unrescued hypo-DMR of Group 2, showing initial intermediate/high DNA methylation levels and H3K4me3-enrichment in WT iPSCs, severe hypomethylation and high H3K4me3 increase in ICF1 iPSCs, which persist in the corrected clones; **(c)** unrescued hypo-DMR of Group 1, showing initial low DNA methylation level and high H3K4me3-enrichment in WT iPSCs, severe hypomethylation and high H3K4me levels in ICF1 iPSCs, which are resistant to correction as in Group 2. Recruitment of methylation-sensitive protein factors with affinity to hypomethylated DNA and highly enriched H3K4me3 regions, and known to compete with DNMTs, such as CTCF or TET enzymes, might contribute to inhibit the binding of the corrected DNMT3B.

The hypomethylated regions associated with the highest abnormal H3K4me3 levels mostly correspond to areas in control iPSCs with active and permissive chromatin features, such as promoters and gene regulatory regions, which are characterized by relatively high H3K4me3 levels in the first place (Fig. 5A). These findings correspond with the observation that centromeric and pericentromeric regions, that retain normal H3K4me3 levels in ICF1 iPSCs despite their hypomethylation, efficiently restore normal DNA methylation profiles in the corrected clones (Fig.1B).

Conversely, the gene-rich distal regions across many chromosomes are characterized by abnormally higher H3K4me3 levels in ICF1 iPSCs, rendering these regions resistant to correction.

We previously showed that the abnormal H3K4me3 levels at subtelomeric regions represent a pre-existing chromatin mark in ICF1 parental fibroblasts [28]. Therefore, the impact of the epigenetic memory on the efficacy of methylation recovery can vary depending on the severity of the initial abnormal epigenome landscape in parental patient fibroblasts. In our study we can associate the degree of epigenetic memory erasure during reprogramming with the residual levels of H3K4 trimethylation in the iPS cell population. This association might explain, at least in part, the variability in DNA methylation recovery that we observe between the corrected clones following gene editing of patient iPSCs.

Additional factors besides H3K4me3 enrichment are predicted to inhibit DNMT3B binding (Fig.6). Interestingly, we found that regions with persisting DNA hypomethylation showed a significant selective enrichment of binding motifs for CTCF and its paralogous BORIS (CTCFL) factors (Fig. 5E). CTCF is categorized as a methylation-sensitive TF, and its occupancy across the genome has been previously reported to be strongly linked with CG methylation [68]. However, recent evidence demonstrated that upon abrogation of DNA methylation, binding of the canonically transcriptional regulator CTCF is largely unaffected and only a limited set of CTCF are methylation sensitive [69, 70]. Therefore, additional factors might be involved in the inhibition of DNMT3B binding and full restoration of DNA methylation.

Alternatively, TET enzymes might be involved in the process of DNA methylation recovery (Fig.6). We showed that DNMT3B acts to preserve the CGI boundaries. When extremely hypomethylated these regions are refractory to methylation regain (Fig 4D and S8A). It has been shown that TET1 specifically accumulates 5-hydroxymethylcytosine at the edges of hypomethylated CGIs, while knockdown of endogenous TET1 induces methylation spreading from methylated edges into hypomethylated CGIs [71]. Moreover, TETs drive global demethylation in the absence of DNMT3s in human ESCs, and thus compete with these enzymes to target many genomic regions [72, 73]. Therefore, it can be hypothesized that they illegitimately bind the DNMT3B targets in ICF1 iPSCs, impeding the recruitment of the corrected DNMT3B protein. Further studies will be necessary to address these possible mechanisms.

Taken together, analysis of remethylation-resistant genomic loci sheds light on the limitations enforced by the abnormal epigenetic landscape of ICF1 chromatin on the ability of the corrected enzyme to carry out its normal activity. This information will assist in designing future strategies for alleviating the obstacles preventing the full recovery of normal DNA methylation patterns in ICF1 cells.

## CONCLUSIONS

In summary, we identified the primary defects in the whole genome DNA methylation landscape of ICF1 syndrome iPSCs caused by DNMT3B loss of function. Remarkably, upon CRISPR/Cas9 correction of the *DNMT3B* mutations, the majority of the hypomethylated sites in ICF1 iPSCs fully restore WT iPSC methylation patterns. However, a small fraction of genomic regions characterized by the lowest DNA methylation levels in ICF1 iPSCs, fail to recover normal DNA methylation levels. Our multi-omic study findings contribute to the identification of molecular factors that influence the ability to correct the ICF1 abnormal epigenome. In perspective, this study emphasizes challenges that will be encountered in correcting inherited epigenetic diseases when gene editing becomes a mainstream therapeutic tool.

## MATERIALS AND METHODS

### Cells and cell culture conditions

In this study we utilized iPSCs from two unrelated ICF1 patients, pR and pG, their isogenic corrected clones (cR7 and cR35, cG13 and CG50) obtained following CRISPR/Cas9 correction of *DNMT3B* mutations, and control UN1-22 iPSCs, defined as WT1 iPSCs [28, 74]. An additional control iPSC line (UWWC1-DS2U from WiCell repository) defined as WT3 iPSCs, was utilized for gene expression qPCR experiments. The iPSC lines were cultured on Geltrex™ LDEV-Free Reduced Growth Factor Basement Membrane Matrix (A1413302, Gibco™) coated plates and maintained in StemMACS™ iPS-Brew XF, human medium (130-104-368, Miltenyi Biotec). Cells were passaged every 4-6 days. Cells were dissociated from plates with StemPro™ Accutase™ Cell Dissociation Reagent (A1110501, Gibco™), and following each passage, 1X of RevitaCell™ Supplement (100X) (A2644501, Gibco™), was added to the media. Lymphoblastoid cell lines (LCLs) used in the study were derived from two unrelated ICF1 patients: ICF1p1 LCL (GM08714, Coriell Cell Repository) and ICF1p2 LCL (PT5 [20]). In addition, we utilized LCLs from one healthy donor (WT LCL, (20)). LCLs were cultured in RPMI1640 Medium, Glutamax™ Supplement (61870036, Gibco™) supplemented with 1X of Penicillin-Streptomycin (15140122, Gibco™) and 15% heat-inactivated fetal bovine serum (ECS5000L, Euroclone). The identity of iPSCs and LCLs was verified by Sanger sequencing of the patient-specific mutations and of the silent mutations incorporated during the editing process into the *DNMT3B* gene. All cells were negative in a screen for mycoplasma.

### Whole Genome Bisulfite Sequencing (WGBS) and data processing

We extracted genomic DNA from WT1 iPSC, ICF1 iPSCs and their respective corrected clones, using the Wizard® Genomic DNA Purification Kit (A1125, Promega). Approximately 100ng of genomic DNA was used for the WGBS experiment. The bisulfite-treated library preparation and the 100bp length paired-end (PE) sequencing were performed at Genomix4Life S.r.l (Salerno, Italy) using the Illumina NextSeq 500 system. After adaptor trimming and quality filtering with Cutadapt [75] we used Bismark [76] to align the PE reads (Table S1) to the bisulfite-converted hg38 reference assembly and extracted the cytosine methylation calls. The methylation level was expressed as the ratio of the number of Cs over the total number of Cs and Ts at base resolution and across the regions of interest. We employed the MethylKit R package to identify the Differentially Methylated Regions (DMR) [77]. Hypomethylated DMRs (hypo-DMRs; 1kb-size) were detected by setting a threshold for the difference in methylation percentage (meth.diff score) > 25%; adjusted p-value (q) < 0.01 and level = “hypo”, when individually comparing ICF1 iPSCs patients, and clones to WT1 iPSCs. Then, we filtered the lists of hypo-DMRs by removing from downstream analyses the hypo-DMRs, where the internal control WT1 iPSCs showed a difference over 0.2 in the methylation ratio compared to WT2, an additional control iPSC corresponding to the human dermal fibroblasts-derived iPSC downloaded from GEO dataset *GSM1385983* [31]. The ICF1 hypo-DMRs were defined as rescued in corrected clones if, compared with the internal control WT1, had i) meth.diff score < 25%, level=”hypo” and/or q > 0.01; or ii) meth.diff > 25% and level=”hyper”.

We annotated the hypo-DMRs to genes using the *annotatePeak* function from the ChIPseeker R package [78] and reported only the nearest gene feature per region. The gene features were classified into promoters (up to 2kb upstream of the transcription start sites (TSS) to 500bp downstream to the TSS), 5’ UTR, Exons, Introns, 3’ UTR, downstream (< 3kb downstream transcription end sites (TES)) and distal intergenic (> 2kb distal from TSS and > 3kb from TES). Genes (ENSGs) annotated to the hypo-DMRs were provided as the input to perform functional enrichment analysis on the PANTHER database web interface [79]. In addition, transcription factor (TF) motif analysis was performed on the ICF hypo-DMRs using HOMER suite [80]. For detailed descriptions and further downstream analyses, see “Supplementary Data”.

### Chromatin Immunoprecipitation Sequencing (ChIP-seq) and data processing

We performed chromatin immunoprecipitation (ChIP) to identify DNMT3B, H3K4me3 and H3K36me3 enriched regions in WT1, ICF1 iPSCs and their corrected clones. 3x10^6^ cells were fixed twice for 40 minutes in 20 mM ethylene glycol bis (succinimidyl succinate) (EGS), and for 10 minutes in 1% formaldehyde solution to promote cross-linking. Excessive formaldehyde was quenched with 125 mM glycine solution and cells were washed twice with PBS. Cells were then lysed in a solution containing 10 mM Tris-HCl pH 8, 100 mM NaCl, 10 mM EDTA, 0.25%, Triton X-100, EDTA and cOmplete™ Protease Inhibitor Cocktail (PICS) (11697498001, Roche). Nuclei were lysed in a solution of 1% SDS containing 50 mM HEPES-KOH (pH 7.5), 150 mM NaCl, 1% SDS, 2 mM EDTA, 1% Triton X-100, 0.1% NaDOC and PICS. After centrifugation, the chromatin pellet was resuspended in the same solution containing 0.1% SDS. For ChIP with the DNMT3B antibody, sonication was conducted using the Covaris S220 apparatus (12 cycles, 40” on, 20” off, intensity 4). Sonicated chromatin was incubated overnight at 4°C with Magna ChIP™ Protein A Magnetic Beads (16661, Merck Millipore) and an anti-DNMT3B antibody (C15410218, Diagenode). Decrosslinking from beads was performed in a 1% SDS solution containing Tris-HCl pH8 and 10 mM EDTA for 2 hours at 69°C with agitation. Decrosslinking of the DNA-protein complex was performed by adding Proteinase K (20 ng/mL) for 2 hours at 62°C with agitation. DNA was then purified using Expin^TM^ PCR SV (103-102, GeneAll®). ChIP with antibodies for histone marks (H3K4me3, ab8580; H3K36me3, ab9050, Abcam) was performed as previously described [81]. ChIP libraries (200-500 bp fragment sizes) were generated from two biological IP replicates, one input for DNMT3B and one pooled input for histone marks per iPSC sample. Single-end (SE) sequencing performed on the Illumina Hi-Seq 2500 system at the Technion Genome Center (Haifa, Israel) produced reads of 100bp length from each IP and input (Table S1). The pre-processing steps included quality analysis and trimming of sequenced SE reads, alignment to the hg38 reference assembly using Bowtie2 aligner [76] and conversion of de- duplicated BAM reads to BED files for peak-calling. We used the SICER2 tool [82] to call enriched IP peaks versus their input and to identify Differentially Enriched Regions (DERs) in ICF1 iPSCs and their corrected clones compared to WT1 iPSCs. Increased H3K4me3 DERs identified in ICF1 iPSCs were defined as rescued in corrected iPSCs if not detected in the list of increased H3K4me3 DERs compared to WT1 or if called as decreased H3K4me3 DERs in this comparison. The rescue of decreased DNMT3B DERs in corrected iPSCs was measured following a similar criterion. Average ChIP-seq enrichment profiles and heatmaps were generated using ngs.plot [83]. For a detailed description and further downstream analyses, see “Supplementary Data”.

### RNA-sequencing and data processing

Total RNA extraction from iPSCs and LCLs was performed using TRI Reagent® (T9424, Sigma-Aldrich), according to the manufacturer’s protocol. RNA was quantified using Nanodrop 1000 Spectrophotometer (ThermoScientific) and the absorbance ratio 260/280 was measured to assess the RNA purity. Extracted RNA was further purified from genomic DNA using the TURBO DNA-*free*™ Kit (AM1907, Invitrogen™). We tested the absence of residual DNA, using RNA as a template to amplify a genomic region (MYOD1 gene). The quality and integrity of RNA was assessed by measuring the RNA Integrity Number (RIN) with the Bioanalyzer instrument (Agilent). Strand-specific libraries were prepared according to Illumina’s instructions and the libraries were sequenced at the IGA (Udine, Italy) using the Illumina Hi-Seq 2500 platform. We obtained PE reads of 125bp length from two technical replicates per each iPSC sample (Table S1). First, we performed quality control to remove low-quality PE reads and adapter sequences and then we aligned the sequences to the hg38 reference assembly using HISAT2 [84] and reverse-strand specific parameters. Next, we quantified gene expression as read counts using the *FeatureCounts* function from Rsubread package [85], and identified the differentially expressed genes (DE) using the NOIseq R package [86], after normalization and batch effect removal. We applied a threshold of posterior probability (pp) > 0.9 to call DE genes and divided them into up- and down-regulated DE genes based on the log2 fold change in expression vs WT1 (+/- log2FC).

For the analyses including the integration of gene expression with DNA methylation (Fig. S6C, Fig. 3A) we relaxed the threshold, which defines genes as DE, to pp > 0.8. DE genes identified in ICF1 iPSCs were defined as fully rescued if their posterior probability in the comparison vs WT1 was pp < 0.5 in both clones, as partially rescued if pp < 0.5 in only in one clone, and as slightly rescued if in both corrected clones 0.5 < pp < 0.8.

### Quantitative Real time PCR

1 µg of RNA derived from iPSCs was reverse-transcribed using 100ng of Random Primers (48190011, Invitrogen™) and 100U of SuperScript™ II Reverse Transcriptase (18064022, Invitrogen™) in a T100™ Thermal Cycler (Biorad), according to manufacturer’s protocol (5’ at 65°C, 2’+10’ at 25°C, 40’ at 42°C and 15’ at 70°C). Real-time quantitative PCR (RT-qPCR) was performed using SsoAdvanced™ universal SYBR® Green supermix (1725270, Bio-Rad) in a Bio-Rad iCycler, according to manufacturer’s protocols. Expression levels were normalized to the *GAPDH* gene by the ΔΔCt method.

Immunoprecipitated samples and corresponding mock samples (negative controls to measure background) were used for ChIP-qPCR. The enrichment of DNA was calculated in terms of % input = 2- ΔCt x 100, where ΔCt (threshold cycle) was determined by CtIP sample - CtInput. Primer sequences for gene expression and ChIP analyses appear in Table S2.

### Statistical analysis

Specific statistical tests performed in genome-wide studies are described in figure legends and the “Results” section. P-values were adjusted with Benjamini-Hochberg method with FDR < 0.01 (BH-FDR) and the significant p-adjusted values were represented as follows: *=p-adjust < 0.01; **=p-adjust < 0.001; ***=p- adjust < 0.0001. The data analyzed using RT-qPCR and ChIP-qPCR is presented as mean ± SD from independent triplicates, each amplified twice. Statistical analyses were performed using a one-tail two-sample Student’s t-test (*=p-value < 0.05, **=p-value < 0.01, ***=p-value < 0.001). In the case of RT-qPCR, statistical significance was calculated comparing each ICF1 or corrected iPSCs to each WT (WT1 and WT3) iPSCs separately and the least significant values were reported in the corresponding Figures.

## AVAILABILITY OF DATA AND MATERIALS

Sequencing data have been deposited in GEO under SuperSeries accession number GSE197925. The ChIP-seq, RNA-seq and WGBS data generated are available under the accession number GSE197922, GSE197923, GSE197924, respectively. The public datasets analyzed in this study are available in the following repositories: The WT2 external control iPSC WGBS dataset (GSM1385983, https://www.ncbi.nlm.nih.gov/geo/query/acc.cgi?acc=GSM1385983) and the methylation array data of primary blood samples from ICF1 patients (GSE95040, https://www.ncbi.nlm.nih.gov/geo/query/acc.cgi?acc=GSE95040). The list of links to all websites of data, software or tools utilized in the present article are available in Supplementary Data.

## ETHICS APPROVAL AND CONSENT TO PARTICIPATE

Not applicable.

## CONSENT FOR PUBLICATION

Not applicable.

## ACKNOWLEDGEMENTS

We are grateful to Miriam Gagliardi for her valuable help in the initial phases of the study and for the critical reading of the manuscript. We thank Dario Righelli and Ankit Verma for their advice about the application of bioinformatic tools for NGS data analysis. We are grateful to Daniel Kornitzer, Floriana Della Ragione and Andrea Riccio for their comments on the manuscript and for helpful discussions.

## AUTHORS’ CONTRIBUTIONS

V.P.K. carried out the computational and statistical analysis of WGBS, ChIP-Seq and RNA-Seq datasets.

B.M. performed the ChIP assays and the experimental validations of the sequencing data. S.T. performed the ChIP assays. M.K. contributed to the RNA-Seq data analysis; M.S., C.A., S.S. and M.R.M. interpreted data. M.R.M., S.S. and C.A. and conceived and coordinated the work and wrote the manuscript. All authors read and commented on the manuscript.

