## Supplemental Data and Figures for "The rescue of epigenomic abnormalities in ICF1 patient iPSCs following *DNMT3B* correction is incomplete at a residual fraction of H3K4me3-enriched regions"

### SUPPLEMENTARY DATA

#### Genome-wide DNA methylation analysis (WGBS)

We assessed the quality of sequenced PE reads using FastQC prior and post-trimming. Using Cutadapt [1], we filtered out the raw read pairs with Phred score < 30, read length < 40 and trimmed the Illumina adapter sequences by applying the following parameters: -u 7; -U 7; -q > 30; -m > 40; -a AGATCGGAAGAG -A AGATCGGAAGAG. Summarized reports of pre- and post-trimmed PE reads were obtained using multiQC software [2]. We used the Bismark program with default parameters for the bisulfite-conversion and indexing of the hg38 reference genome (canonical chromosomes including chr 1-22; X and Y), alignment of PE reads to the reference index, and the removal of PCR duplicates. We obtained the per-base cytosine methylation report (CG, CHG, CHH) using *methylation extractor* command (parameters: -p --no_overlap --bedGraph --counts --zero_based –cutoff 2 --cytosine_report –CX_context). We evaluated the methylation level of cytosines (Cs) from both strands in the context of CG, whereas we measured the methylation level of strand-specific Cs for the non-CGs. We tiled the genome into 1kb windows with at least 2 Cs and minimum coverage of 5 reads per tile and identified the Differentially Methylated Regions (DMRs) using the MethylKit R package (meth.diff score>25%, qvalue=0.01) and selected the hypo-DMRs with type="hypo" where the meth.diff score is the difference between the methylation percentages.

To filter the hypo-DMRs in the comparison between ICF1 and WT1 iPSCs, we downloaded the single-end (SE) FASTQ file of the additional iPSC control WT2 (see Methods) using the fasterq-dump tool from SRA Toolkit [3] and aligned them to the hg38 reference genome using the Bismark aligner with default parameters for SE mapping. We performed the extraction of methylation content, coverage calculation for WT2. Then, we filtered out the ICF1 hypo-DMRs when we observed a difference > 0.2 between the methylation levels (expressed as the ratio of the number of Cs over the total number of Cs and Ts) in WT1 and WT2 iPSCs, as described in Methods (Whole Genome Bisulfite Sequencing and data processing). The hypo-DMRs identified in chrY were removed from pR and its corrected clones. The ICF1 hypo-DMRs were defined as rescued in the corrected clones if compared with the internal control WT1 they had i) meth.diff score < 25%, level="hypo" and/or q > 0.01; or ii) meth.diff > 25% and level="hyper".

Using the *annotatePeak* function from the ChIPseeker R package, we annotated the DMRs to single genomic features (parameters: tssRegion=c(-2000,500), annoDB="org.Hs.eg.db", overlap="all", genomicAnnotationPriority=default) based on the criteria described in the Methods (Whole Genome Bisulfite Sequencing and data processing).

We performed the functional enrichment analysis of the genes annotated to promoter and gene body hypo-DMRs independently using PANTHER applying the settings: Analysis- Statistical overrepresentation test, Annotation set- PANTHER GO-Slim Biological Processes (GO-BP), Background: Homo sapiens genes, Test type- Fisher's exact test, Correction- False Discovery Rate (default FDR < 0.05).

We assessed the statistical significance of the overlap between ICF1 hypo-DMRs and the Regions of Interest (ROIs) (i.e., CGI, GH, DMV, hypo-DMRs LCLs, ChIP-seq DERs) by shuffling using the

*enrichPeakOverlap* function from ChIPseeker with the following parameters: TxDb=hg38_ensembl, pAdjustMethod="BH", nShuffle=4000.
To compare our results with the microarray-based profiles of DNA methylation in ICF1 patients' whole blood ([4], we downloaded the methylation arrays from GSE95040, calculated the median beta ($\beta$) values (i.e., the methylation level expressed in the [0,1] range) of the control samples ($\beta_{Control}$) and ICF1 samples ($\beta_{ICF1}$), and identified the hypomethylated regions in ICF1 blood as those in which $\beta_{ICF1} - \beta_{Control} < -0.2$.
We divided the ICF1 hypo-DMRs into four groups using k-means clustering based on the mean methylation levels across the iPSCs samples. Group1 consisted of hypo-DMRs with the lowest mean methylation level, while Group 4 had the highest mean methylation level across all iPSCs analyzed. Group 1 and 2 consisted of hypo-DMRs showing lower methylation level in ICF1 iPSC while Group 3 and 4 exhibited relatively higher ICF1 methylation level along with varying degrees of methylation rescue in corrected clones.
For Transcription Factor (TF) motif enrichment analysis, first, we selected the subsets of the hypo-DMRs overlapping with decreased DNMT3B DERs in pR and pG, respectively. Next, the subsets of hypo-DMRs from Group 1 and 2 (which showed low methylation recovery in corrected iPSCs) were combined. Similarly, the subsets of hypo-DMRs belonging to Group 3 and 4 were combined. Then, we performed motif enrichment analysis using *findMotifsGenome.pl* from the HOMER suite (parameters: --size given and genome "hg38"), on i) the combined subset from Group 1 and 2, and ii) the combined subset from Group 3 and 4. In addition, we also performed known motif enrichment analysis on all the hypo-DMRs overlapping with decreased DNMT3B DERs (DNMT3B-Dec) in ICF1 iPSCs to identify TFs regulating DNMT3B-mediated methylation at genomic target regions.

*Data visualization*

We used the ViewBS toolkit [5] for obtaining genome-wide methylation coverage (*GlobalMethLev*) for each cytosine context (Fig. S1A-C) across all iPSC lines. We generated and visualized the profile-plots of the average weighted methylation level (*MethOverRegion)*, the distribution of methylation levels and summary statistics (*MethHeatmap*) for specific ROIs using the ggplot2 and pheatmap R packages (colorRampPalette(colors=c("yellow", "blue"))(25), cluster_cols=FALSE, cluster_rows=FALSE). Heatmap representation of methylation levels across ICF1 hypo-DMRs and their rescue status (Fig. 1A) was generated using the ComplexHeatmap R package. The ICF1 hypo-DMRs were first clustered (k-means) into four groups, then further defined based on their rescue status: "full rescue" (rescued in both corrected clones), "partial rescue" (rescued in only one corrected clone) and "no rescue" (remaining hypo-DMR in both corrected clones). The integrated heatmaps was constructed using the *heatmap* function with the following parameters: split=k-means_row_order; col (methylation level)=colorRamp2(c(0,25), c("yellow","blue")) or col (rescue status)=c("green3", "orange", "red).
We plotted the chromosomal distribution (22 autosomes; Fig. 1B) of pR and pG hypo-DMRs, rescued hypo-DMRs and CpG islands (CGI) using the genomicDensity function from the R circlize package (6)

and represented as density line plots along with each chromosome ideogram using the gtrellis_layout function from the gtrellis R package (7). We set the window.size parameter to 1Mb for hypo-DMRs and 1kb for CGI, respectively.

We represented the genomic distribution of the annotated hypo-DMRs and those rescued in corrected clones by gradient donut charts (Fig. 2A) generated with the *patternpie* and *patternring1* functions from the patternplot R package.

We visualized the enriched Biological Processes (GO-BP) associated with the pR and pG common hypo-DMR annotated genes (ENSGs) as a Multidimensional Scaling (MDS) scatter-plot (Fig. 2B) by modifying the color scale (-log10 of the adjusted p-value of the GO term) and dot size (log2 of the number of hypo-DMR associated genes in GO term) in the Rscript output from the REVIGO online tool [6]. In addition, we represented the proportion of the pR and pG common hypo-DMRs rescued in genes associated with enriched GO terms as a stacked barplot (Fig. 2C) using the *geom_bar* function from ggplot2.

We obtained the Upset plot distribution of the ICF1 hypo-DMRs associated with regulatory regions (CGI/GH) in ICF1 iPSCs and in corrected iPSC clones (Fig. 4D and S8C) using the UpsetR package with the following parameters: order.by="freq", keep.order="TRUE", group.by="degree", decreasing="TRUE". Boxplots indicating the methylation levels across rescued and unrescued CGI and GH hypo-DMRs were generated using ggplot2.

We quantified the overlap of hypo-DMRs with the closest ROI using the *subsetByOverlaps* from the GenomicRanges R package. Next, we used the Venn diagram to depict the overlaps through the *draw.pairwise.venn* function from the VennDiagram R package [7]. Then, we computed the shortest distance between the hypo-DMRs and ROIs using the *annotatePeakInBatch* from ChIPpeakAnno with output set to "nearestLocation" and plotted the distribution of the shortest distances using the *geom_hist* function from ggplot2.

We converted the bedgraph files from Bismark methylation caller to BigWig tracks using *bedGraphtoBigWig* utility and hosted them at Cyverse Discovery Environment [8]. The methylation coverage files across the genome were visualized on the UCSC genome browser [9] with the following settings: track type=bigWig, visibility=full, viewLimits=default, windowingFunction=mean, smoothingWindow=10, color=100,0,0). The filtered hypo-DMRs were uploaded as BED tracks with differential methylation score indicated by gradient grey-scale boxes. Four shades of grey ranging from light grey to dark grey correspond to -25 to -39, -40 to -59, -60 to -79, and -80 to -100 hypo-DMRs score.

We visually compared the hypo-DMRs obtained in ICF1-iPSCs to hypomethylated regions (HMR) in wild-type human embryonic stem cells (hESCs) and hESCs in which DNMT3B was knocked-out and knocked-down available in UCSC genome browser database. HMR is defined as regions across the genome with low-methylation levels identified using the MethPipe package [10]. We displayed the following HMRs available in MethBase track hub: three wild-type hESCs - H1 [11]; H9 [12]; HUES6 [13], two DNMT3B knock-out hESCs - early (2–7) and late (17–22) passage [14] (3BKO) and one shRNA DNMT3B knock-down hESCs [12] (3BKD).

**ChIP-seq data analysis**

We assessed the quality of the sequenced single-end (SE) reads from IP and input using FastQC prior and post-trimming. Using cutadapt, we filtered and adaptor-trimmed the SE reads using the following parameters: -q > 30; -m > 40; -a AGATCGGAAGAG. We aligned the retained reads to the custom hg38 reference genome with canonical chromosomes only (GRCh38/hg38 primary assembly) using Bowtie2 with the default parameters. Furthermore, we filtered out aligned SE reads with MAPQ score < 20 using the *samtools view* -q 20 setting [15], then removed PCR duplicates using the *samtools markdup* command.

We identified the narrow H3K4me3 peaks (enriched regions) and differentially enriched regions (DERs) for each IP replicate using the *sicer_df* command (SICER2) setting the following arguments: -f 200, -fdr 0.00001, -fdr_df 0.01, -egf 0.88, -w 200, -g 200, -s hg38. For broad H3K36me3 peaks (enriched regions), we increased the gap size to -g 1400 (optimized by plotting a curve with the sum of island counts at different gap sizes). Since DNMT3B peak structure reflects H3K36me3 peak enrichment, we also set -g to 1400 with -fdr and -fdr_df set to 0.0001. After that, we removed the enriched peaks and DERs overlapping the ENCODE hg38 blacklist regions [16] with the *intersectBed* function from the BEDtools suite [17]. We defined the consensus peaks and DERs as those overlapping at least 1bp in both IP replicates. DERs with contradictory fold change (FC) direction (increased or decreased) between IP replicates were removed. We also defined the increased H3K4me3 DERs identified in ICF1 iPSCs as rescued in corrected iPSCs if not detected in the list of increased H3K4me3 DERs compared to WT1 or if called as decreased H3K4me3 DERs in this comparison. We used the same criterion to define the rescue of decreased DNMT3B DERs in corrected iPSCs.

To calculate the enrichment of an IP across ROIs, we first counted the reads in both IP and input using the *featurecounts* function with the following settings: annotation=custom SAF file, useMetaFeatures=FALSE, allowMultiOverlap=FALSE, strandSpecific=0, CountMultipleMappingReads=TRUE). Then, we performed batch-effect removal using the *ArySyNseq* function (factor="run", batch=TRUE, norm="n") to remove the bias likely introduced by the different sequencing runs. Subsequently, we normalized the batch-corrected read counts by the ChIP-seq library size and computed the fold change (FC) = Normalized read count of IP/Normalized read count of input (IP/input).

We first considered the DNMT3B peaks in WT1 and then computed the FC of these peaks across all the iPSCs. Next, we calculated the FC ratio of WT1 to pR/pG and identified WT1 peaks showing lower DNMT3B enrichment in ICF1 iPSCs by applying a threshold of FC ratio > 1.2. From these peaks, we further subsetted those showing higher enrichment (FC) scores in corrected clones compared to ICF1 to compute the rescue percentage described in the results.

*Data visualization*

We generated the hybrid plots (Dot, box and violin plot; Fig. S3A) to represent the fold enrichment of DNMT3B in pR-related (pR, cR7, cR35) and pG-related (pG, cG13, cG50) samples using the ggstatsplot R package [18] with the following parameters: plot.type="boxviolin", type="nonparametric", p.adjust.method="BH", centrality.type="parametric" (to denote the mean enrichment score). Then, we performed two-sample, two-sided paired Wilcoxon test to compare WT1 DNMT3B peaks fold enrichment (FC) to the other samples in the plot using the *Stat_compare_means* function from the ggpubr R package (method="wilcox.test", paired=TRUE and p.adjust.methods="fdr"). We denoted the p-adjusted values for each comparison as follows: *p-adjust < 0.01; **p-adjust < 0.001; ***p-adjust < 0.0001.

We plotted the chromosomal distribution (22 autosomes; Fig. S3B) of pR and pG hypo-DMRs, CpG islands (CGI) as density line plots alongside the rainfall plots representing the fold change of pR- and pG-set samples at pR and pG vs WT decreased DNMT3B DERs respectively using the gtrellis_layout function from the gtrellis R package (7). We set the window.size parameter to 2Mb for hypo-DMRs and 1kb for CGI respectively. Following the TF motif enrichment analysis at ICF1 hypo-DMRs intersecting DNMT3B decreased DERs, the TF binding at the hypo-DMRs was confirmed by calculating the significance of the overlap between the hypo-DMRs and the TF ChIP-seq enriched peaks (ENCODE Accession number for E2A: ENCFF658WIO; EBF1: ENCFF249SVT; c-MYC: ENCFF700CXD derived from GM12878 LCLs and SCL/TAL1: GSM1816082 derived from Fetal HSPCs) and represented as Venn diagrams.

We generated the multi-omics integrated heatmaps (Fig. 5A) displaying the DNA methylation levels, histone marks and DNMT3B enrichment, genomic annotation, and the rescue status of CGI-GH hypo-DMRs in pR and pG- related iPSCs using the *heatmap* function from the ComplexHeatmap R package. We divided the genomic features annotated to the hypo-DMRs into three groups: "Promoter", "Gene body" (Intron/Exon/5'UTR/3'UTR) and "Distal intergenic". The hypo-DMRs were divided into two groups based on their rescue status: "full rescue" (rescued in both corrected clones) and "no rescue" (remaining hypo-DMR in both corrected clones). The rows of the heatmap were clustered (k-means) into four groups, similar to the heatmap in Fig.1A. The color scale for methylation levels and ChIP-seq enrichment (FC) was produced using the *RcolorBrewer* package.

We plotted the H3K4me3 fold enrichment (FC) density plot (Fig. 5B) at H3K4me3 DERs overlapping pR and pG hypo-DMRs using the geom_density_ridges function from the ggridges R package. For constructing the density plot, we calculated the H3K4me3 FC at the ROIs across pR-related and pG-related iPSCs and grouped them into three categories based on the ranking of FC from lowest to highest in pR/pG ICF1 iPSCs.

We obtained the bigWig coverage tracks of ChIP-seq replicates and input by converting BAM to BED files, extending the read length by 100bp (fragment size ~ 200bp) using *slopBed* and computing the read coverage using the *genomecov* function in the BEDtools suite. We normalized the bedGraph tracks to the library size, sorted and converted to bigWig files, and hosted them at Cyverse Discovery Environment and uploaded on a UCSC genome browser session with the following setting: (track type=bigWig, visibility=full,viewLimits=default, windowingFunction=mean, smoothingWindow=10, color=0,0,255[DNMT3B]/ 128,0,128[H3K36me3]/ 0,100,0[H3K4me3]). The enriched consensus peaks and DERs were also uploaded as bedGraph tracks.

**RNA-seq data analysis**

We sequenced the pR-related and the pG-related samples in two separate batches, with the WT1 replicates sequenced in both batches. First, we filtered out the low-quality reads and trimmed the adapters in the strand-specific PE reads using Cutadapt by setting the following parameters: -q 30 -m 40 -a AGATCGGAAGAG -A AGATCGGAAGAG. Then, we aligned the trimmed reads to the hg38 reference genome (canonical chromosomes only) using HISAT2 with the following options: -p 8 --dta --rna-strandness RF.

Next, we quantified the gene expression by counting the reads mapping to genes (ENSGs) using the *featurecounts* function from the Rsubread package with the following settings (annot.ext=”hg38.v85" gtf file, useMetaFeatures=TRUE, allowMultiOverlap=FALSE, strandSpecific=2, CountMultiMappingReads=FALSE). After that, we filtered out zero count or low count genes (i.e., those with CPM < 0.5) using the Proportion Test Method from the NOIseq R package. Overall, we obtained a list of expressed genes (n=18,077) excluding the genes expressed from the chromosome Y since the iPSCs were derived from individuals of different biological sex (pR- female; pG-male; WT1- male). Then, we normalized the raw counts across the samples using the Upper Quartile (UQUA) method.

The inspection of the Principal Component Analysis (PCA) on the normalized read count matrix revealed a batch effect between the samples due to the two different sequencing runs. Therefore, we performed the batch-effect removal using the *ArySyNseq* function with parameters: factor="run", batch=TRUE, norm="n", where the WT1 counts from the two runs were henceforth considered as one control sample with four replicates while every other sample (ICF1 and corrected iPSCs) had two replicates. After that, we identified the differentially expressed (DE) genes using the *noiseq* function from the NOIseq package. We compared the expression of ICF1 iPSCs and corrected iPSCs versus WT1 iPSCs, and we defined the DE genes in each comparison as those genes with a posterior probability (pp) > 0.9. As a further quality control of the gene expression profile in our WT1 iPSCs, we downloaded the RNA-seq FASTQ reads of additional WT iPSCs [19] [20] and ESCs [21] and processed them as described above to obtain the Log2 of the UQUA normalized counts (i.e., log2.UQUA external control). The DE genes in the comparisons of pR and pG vs WT1 were filtered out by removing those genes with abs(log2.UQUA WT1 - log2.UQUA external control) > 2 and showing a fold change with opposite sign when compared to patient iPSCs.

*Data visualization*

Scatterplots showing the log10 normalized counts (Fig. S6A) or the log2FC of the expressed genes (Fig. S6C) were generated using the ggplot2 package. We used Upset plots (Fig. S6B) to visualize the distribution of DE genes in pR and pG ICF1 iPSCs and their differential expression status in the corrected iPSCs vs WT1 using the UpsetR package (order.by="freq", keep.order="TRUE", group.by="degree", decreasing="TRUE"). We depicted the changes in the expression level of DE genes (pp > 0.8) associated with ICF1 hypo-DMRs and annotated to specific gene features (promoter, exon, intron, 3'UTR) as boxplots (Fig. 3A) using the *geom_boxplot* function (ggplot2). We grouped the DE genes in ICF1 and their corrected clones vs WT1 in each gene feature category. Then, we performed a pairwise comparison (two-sample, two-sided, paired Wilcoxon test) between the log2FC of the ICF1 vs WT1 genes associated with promoter hypo-DMRs and those annotated to other genomic feature categories.

The genes expressed in WT1 were ranked by their log2 FPKM normalized counts and divided into four quartiles with Q1 and Q4 denoting the lowest and highest expressed genes in WT1 iPSCs, respectively. The ChIP-seq enrichment of WT1 for the entire list of ranked expressed genes was represented using a heatmap, while the average enrichment across each quartile was visualized as a profile plot (Fig. S9A). We used the *ngs.plot.r* command with following settings: "-G hg38, -R gene body (DNMT3B/H3K36me3) or TSS (H3K4me3), -E custom_gene_list, -GO none, -YAS 0,0.10/0.15/1.5 (DNMT3B/H3K36me3/H3K4me3), -L 2000, -LEG 1, -SE 0" to obtain the heatmaps and profile plots.

Aligned BAM files of each iPSC line were sorted by the chromosome position, converted to bedGraph, and bigWig files and then hosted at Cyverse Discovery Environment. The bigWig files were visualized as coverage tracks on UCSC genome browser using the following setting: (tracktype=bigWig, viewLimits=0:1, windowingFunction=mean, smoothingWindow=10, visibility=full).

**Reference genome assembly for sequencing alignment and annotation**

We used the GRCh38/hg38 human reference genome assembly (ftp://ftp.ensembl.org/pub/release102/fasta/homo_sapiens/dna/Homo_sapiens.GRCh38.dna_sm.primary_assembly.fa.gz) for the sequence alignment performed in both epigenomic and transcriptomic analyses. For transcriptome annotation, we used the *Homo_sapiens.GRCh38.85_canon_chr_header.gtf* (a filtered version of GRCh38/hg38.v85 annotation file that includes only canonical chromosomes) downloaded from the Ensembl FTP server.

The hg38 chromosome sizes were downloaded via the UCSC Genome Browser FTP server.

We downloaded satellite repeat regions from the RepBase Update database. We downloaded the CpG islands (cpgislandExt), Genehancer regulatory elements, and the ENCODE cCREs (hg38) tracks using the UCSC table browser. We obtained the DNA-methylation valleys (DMVs) BED file from [22]. For the DMRs annotation to the genomic features, we built the TxDb object from the hg38 Ensembl database using the *makeTxDbFromEnsembl* function in the *ChIPseeker* R package. The hg19 tracks downloaded from public databases were converted to hg38 using the *LiftOver* command-line tool.

**List of website links of data, software and tools utilized in the present article:**

| Sofware/Package | Version | Source |
|---|---|---|
| **BEDtools** | v2.29.2 | https://github.com/arq5x/bedtools2 |
| **Bismark** | v0.19.1 | https://github.com/FelixKrueger/Bismark |
| **Bowtie2** | v2.3.4.3 | https://github.com/BenLangmead/bowtie2 |
| **ChIPseeker** | v1.29.1 | https://bioconductor.org/packages/ChIPseeker/ |
| **ChIPpeakAnno** | v3.27.6 | https://bioconductor.org/packages/release/bioc/html/ChIPpeakAnno.html |
| **ComplexHeatmap** | v2.9.4 | https://bioconductor.org/packages/ComplexHeatmap/ |
| **Circlize** | v0.4.13 | https://cran.r-project.org/package=circlize |
| **Cutadapt** | v1.9.1 | https://pypi.org/project/cutadapt/1.9.1/ |
| **DeepTools** | v3.3.2 | https://github.com/deeptools/deepTools |
| **FastQC** | v0.11.5 | https://www.bioinformatics.babraham.ac.uk/projects/fastqc/ |
| **GenomicRanges** | v1.45.2 | https://bioconductor.org/packages/release/bioc/html/GenomicRanges.html |
| **gtrellis** | v1.22.0 | https://bioconductor.org/packages/release/bioc/html/gtrellis.html |
| **ggplot2** | v3.3.3 | https://cran.r-project.org/package=ggplot2 |
| **ggfortify** | v0.4.12 | https://cran.r-project.org/package=ggfortify |
| **ggridges** | v0.5.3 | https://github.com/cran/ggridges |
| **ggstatsplot** | v0.8.0 | https://indrajeetpatil.github.io/ggstatsplot/ |
| **GRCh38/hg38 GTF** | v85 | http://ftp.ensembl.org/pub/release85/gtf/homo_sapiens/Homo_sapiens.GRCh38.85.chr.gtf.gz |
| **GRCh38/hg38 chromosome sizes** | | http://hgdownload.cse.ucsc.edu/goldenpath/hg38/bigZips/hg38.chrom.sizes |
| **HISAT2** | v2.1.0 | https://github.com/DaehwanKimLab/hisat2 |
| **HOMER** | v4.11 | http://homer.ucsd.edu/homer/ |
| **methylKit** | v1.16.0 | https://bioconductor.org/packages/release/bioc/html/methylKit.html |
| **MultiQC** | v1.9 | https://pypi.org/project/multiqc/1.9/ |

| | | |
|---|---|---|
| **ngs.plot** | v2.63 | https://github.com/shenlab-sinai/ngsplot |
| **NOIseq** | v2.34.0 | https://bioconductor.org/packages/release/bioc/html/NOISeq.html |
| **PANTHER** | v16.0 | http://www.pantherdb.org/ |
| **pathfindR** | v1.6.2 | https://cran.r-project.org/package=pathfindR |
| **patternplot** | v1.0.0 | https://cran.r-project.org/package=patternplot |
| **pheatmap** | v1.0.12 | https://cran.r-project.org/package=pheatmap |
| **plotrix** | v3.8-2 | https://cran.r-project.org/package=plotrix |
| **primer-blast** | | https://www.ncbi.nlm.nih.gov/tools/primer-blast/ |
| **Python** | v.2.7/3.7.6 | https://www.python.org/downloads/release/python-376/ |
| **R** | v3.6.3 | https://cran.r-project.org/bin/linux/ubuntu/#install-r |
| **Rstudio** | v1.2.1335 | https://www.rstudio.com/products/rstudio/release-notes/rstudio-1-2/ |
| **ReviGO** | | http://revigo.irb.hr/ |
| **RepBase** | | https://www.girinst.org/server/RepBase/ |
| **Rsubread** | v2.7.3 | https://bioconductor.org/packages/release/bioc/html/Rsubread.html |
| **SICER2** | v2.0 | https://zanglab.github.io/SICER2/ |
| **SRAToolKit** | v2.10.4 | https://github.com/ncbi/sra-tools |
| **upsetR** | v1.4.0 | https://cran.r-project.org/package=UpSetR |
| **VennDiagram** | v1.6.20 | https://cran.r-project.org/package=VennDiagram |
| **ViewBS** | v0.1.11 | https://github.com/xie186/ViewBS |

### SUPPLEMENTARY FIGURES AND TABLES

***Figure S1***

**CG, CHG and CHH global methylation**

(**A**) (**B**) Average weighted global methylation levels of cytosines as determined by WGBS analysis (weighted based on region size and number of CGs and CHs) at the context of CG, CHG, and CHH (H=A/G/T), in WT1, ICF1, and corrected iPSCs, cR7, cR35, cG13 and cG50. The X-axis denotes the methylation level expressed as the ratio of the number of Cs over the total number of Cs and Ts. The Y-axis indicates the cytosine context.

(**C***)* Average weighted global methylation levels of mCA, mCC, and mCT at the context of CHH and CHG, expressed as the ratio of the number of Cs over the total number of Cs and Ts in WT1, ICF1, and corrected iPSCs.

(**D**) Distribution of the distances between individual hypo-DMRs in pR and pG iPSCs. Each bin in the X-axis denotes the shortest distance (bp) between the compared regions, while the Y-axis indicates the percentage of hypo-DMRs in each bin of 5kb size.

***Figure S2***

**Hypo-DMRs and their rescue at large genomic domains**

**(A-C)** Genome-browser views of large genomic regions including the *IGK* gene cluster at pericentromeric region of chromosome 2 **(A)**, the *IGH* gene locus located at the distal region of chromosome 14 (q32.33) **(B)**, and the long *PTPRN2* gene at the distal region of chromosome 7 (**C**), which demonstrate diffused hypomethylation in ICF1 iPSCs and partial to full recovery of normal DNA methylation level in corrected iPSCs clones. Grey boxes represent hypo-DMRs detected in pR, cR7, cR35, pG, cG13 and cG50 compared to WT1 (internal control) and WT2 (external control). Four shades of grey from light grey to black indicate differential methylation scores compared to WT1 as follows: -25 to -39, -40 to -59, -60 to -79 and -80 to -100.

The six tracks at the bottom illustrate the hypomethylated regions (HMRs; [10]) in WT human embryonic stem cells (hESCs) from H1, HUES9 and H9 lines, followed by DNMT3B-KO hESCs (early and late passage DNMT3B-KO; 3BKO) [48] and shRNA DNMT3B-KD hESCs [47] (3BKD). Increased size of hypomethylation regions is evident in hESCs when DNMT3B is depleted, compared to WT hESCs tracks. The position of genomic sites affected by 3BKO and 3BKD, that are direct DNMT3B targets during *de novo* methylation, overlaps many of the hypo-DMRs in ICF1 iPSCs.

***Figure S3***

**DNMT3B binding enrichment in iCF1 iPSCs and corrected counterparts**

(**A**) Hybrid plots (boxplots and dotplots) showing the fold enrichment of endogenous DNMT3B binding at its target regions (n=19,706 binding sites observed in WT1) obtained by ChIP-Seq analysis in iPSCs of WT1, ICF1 iPSCs and their corrected clones. FC indicates the fold change at DNMT3B peaks over input.

Mean for each sample is indicated by the red dot and adjacent number, while the black line represents the median. Statistically significance differences in DNMT3B enrichment level (FC) between WT1 and other iPSCs were calculated using the non-parametric two-sample, two-sided paired Wilcoxon test Benjamini-Hochberg with False Discovery Ratio (BH-FDR) correction (***=FDR < 0.0001).

**(B)** Genomic density line plots of hypo-DMRs in ICF1 pR and pG iPSCs and rainfall plots of significantly decreased DNMT3B binding regions (DNMT3B-Dec) in ICF1 pR and pG iPSCs and their respective corrected clones. The 22 autosomes are depicted as ideograms in the X-axis with the red band denoting the centromere position. For each chromosome from top to bottom: pR and pG hypo-DMRs line plots, rainfall plots of pR, cR7, cR35 and pG, cG13, cG50 iPSCs at DNMT3B-Dec and CGI distribution (green). The Y-axis of the line plots represents the density of hypo-DMRs defined as the proportion of the regions of interest present in each defined genomic window, while the Y-axis of rainfall plots denote the FC (IP/input) at DNMT3B-Dec. Hypo-DMRs and CGI are partitioned into genomic windows of 2Mb and 1Kb respectively.

***Figure S4***

**Distance between hypo-DMRs and regions with decreased DNMT3B binding**

(**A**) Distribution of distances between hypo-DMRs and regions with significantly decreased DNMT3B binding (DNMT3B-Dec) in ICF1 iPSCs. In each plot, the X-axis denotes the shortest distance (bp) between the compared regions, while the Y-axis indicates the percentage of regions with decreased DNMT3B binding in each bin of 5kb size.

(**B**) A genome browser view of the distance between hypomethylated genomic regions and DNMT3B binding peaks at a selected genomic region of chromosome 19 in WT iPSCs compared with ICF1 iPSCs and corrected counterparts. Dark red tracks display methylation coverage measured by WGBS in WT1 and WT2, ICF1 and corrected iPSCs. CGIs are shown in green. The grey boxes represent the hypo-DMRs in ICF1 iPSCs and their respective corrected clones, as described in Fig. S2.
GeneHancer (GH) regulatory regions (promoters and enhancers, red and grey, respectively) and their long-range interactions are indicated (curved lines). The underneath blue tracks illustrate the hypomethylated regions (HMRs) in WT human embryonic stem cells (hESCs) from H1, HUES9 and H9 lines, followed by DNMT3B-KO hESCs (early and late passage DNMT3B-KO; 3BKO) and shRNA DNMT3B-KD hESCs; 3BKD). Black coverage tracks correspond to DNMT3B binding profiles in iPSCs obtained by ChIP-seq and the grey boxes beneath denote the enriched peaks of DNMT3B binding, with the level of grey corresponding to the fold change of ChIP-seq enrichment.

***Figure S5***

**Hypo-DMRs in ICF1 iPSCs and ICF1 LCLs compared to relative controls**

(**A**,**B**) Venn diagrams representing the intersection of hypo-DMRs in pR and pG iPSCs with (**A**) hypo-DMRs in ICF1p1 LCL compared to WT LCL, obtained from RRBS and WGBS experiments, and ICF1p2

LCL, obtained from RRBS experiment (21), (**B**) hypo-DMRs in ICF1 patient whole blood compared to WT whole blood (ICF1 WB) (39). Statistical significance of the overlap was calculated using the shuffle method. **=p < 0.001.

(**C**) A genome browser view of the *IGHA1*, *IGHG1* and *IGHG3* loci showing hypomethylation in ICF1 iPSCs and the level of methylation recovery in the corrected clones. Dark red tracks display methylation coverage measured by WGBS. Grey boxes below the CGI (green) represent hypo-DMRs detected in patients and corrected iPSCs. The grey gradient of the hypo-DMR boxes indicates the range of differential methylation score, with the darkest indicating the highest, as described in Fig. S2. The underneath blue tracks illustrate the hypomethylated regions (HMRs) in WT human embryonic stem cells (hESCs) from H1, HUES9 and H9 lines, followed by DNMT3B-KO hESCs (early and late passage DNMT3B-KO; 3BKO) and shRNA DNMT3B-KD hESCs; 3BKD).

Brown horizontal boxes represent the corresponding hypo-DMRs in ICF1 LCLs when compared to WT LCL. The layered H3K27Ac, H3K4me1 enrichment and the DNaseI hypersensitive sites (HS) tracks from ENCODE indicate the complex 3' Regulatory Region (3'RR) located downstream to the constant *IGHA1* gene. The position of switch regions (S) and isotype promoters (I) important for initiating the class switch recombination (CSR) are shown at the bottom.

**(D)** A genome browser view of the hypomethylation pattern at *EBF4* and *SMRS* genes identified as part of the immune-related processes enriched in the GO-BP analysis. Hypo-DMRs and HMRs are depicted as described above.

(**E**) Venn diagrams representing the intersection of ICF1 hypo-DMRs with enriched target regions of SCL and c-MYC transcription factors (TFs) playing key roles in hematopoietic fate determination. TFs enriched peaks were obtained from published ChIP-seq datasets and ENCODE [23]. Statistical significance of the overlap was calculated using the shuffle method. **=p < 0.001

(**F**) Venn diagrams representing the intersection of ICF1 hypo-DMRs with regions containing GeneHancer regulatory elements (extended by ±3kb). Statistical significance of the overlap was calculated using the shuffle method. **=p < 0.001

(**G**) Distribution of the distances between hypo-DMRs in pR and pG iPSCs and Gene-Hancer promoters/enhancers. In each plot, the X-axis denotes the shortest distance (bp) between the compared regions, while the Y-axis indicates the percentage of hypo-DMRs in each bin of 3kb size.

***Figure S6***

**Transcriptomic analysis in ICF1 and corrected iPSCs and correlation with DNA hypomethylation**

(**A**) Scatterplots of gene expression levels depicted as log10 of UQUA (Upper Quartile) normalized counts in pR (left) and pG (right) iPSCs (Y-axis) compared to WT1 iPSCs (X-axis). The differentially expressed (DE) genes with posterior probability (pp) > 0.9 in ICF1 compared to WT1 iPSCs are displayed in red (upregulated) and green (downregulated), while the statistically non-significant genes are shown in grey. The DE genes in both or in either pR and pG iPSCs are denoted as diamonds and circles, respectively.

(**B**) Upset plots showing the distribution of DE genes (pp > 0.9) in pR (left) and pG (right) iPSCs and their corrected counterparts. Vertical bars represent the intersection size of DE genes between the depicted iPSCs. Black connected dots in the bottom panel represent DE genes present in each intersection. In details, the black bar indicates genes DE in ICF1 iPSCs and both their corrected clones compared to WT, whereas the second bar to the left includes genes DE only in patient iPSCs and showing full (light green; pp < 0.5 in both clones), partial (medium green; pp < 0.5 in only one clone) or slight (dark green; 0.5 < pp < 0.8 in both clones) rescue in their respective corrected clones. Light grey represents genes with pp > 0.8 in one clone and 0.5 < pp < 0.8 in the second clone, while dark grey denotes genes with pp > 0.8 in both corrected clones. The last two bars represent genes DE in pR or pG and in one of the two corrected clones, with the second clone being partially or slightly rescued (medium green and dark green).
(**C**) Scatterplots describing distribution of genes annotated to hypo-DMR associated regulatory elements in pR (n=3181) and pG iPSCs (n=3319). The X-axis denotes the differential methylation score (expressed as the difference in the methylation percentages) of the hypo-DMRs associated with each gene (average in the case of multiple hypo-DMRs in one gene) and the Y-axis denotes the log2 fold change of the expressed genes. The genes common to both patient iPSCs or unique to pR/pG iPSCs are indicated as diamonds and circles, respectively. Up-regulated and down-regulated genes (pp > 0.9) are depicted in red and green respectively.
(**D**) RT-qPCR of hypomethylated genes *RUNX1, IGHG1* and *LINC00221* in WT iPSCs, ICF1 and corrected iPSCs, WT LCL, ICF1p1 LCL and ICF1p2 LCL. The expression of these genes is unaffected in ICF1 iPSCs but significantly altered in ICF1 LCLs when compared to WT LCLs. Each bar represents the mean of the relative expression compared to the expression of *GAPDH* in the same sample. Error bars represent SEM of at least three experimental repeats. Primer sets used for RT-qPCR analysis are reported in Table S2.
(**E**) RT-qPCR of hypomethylated genes *CD74* and *PLA2G4C* whose expression is deregulated in both ICF1 iPSCs and in ICF1 LCLs when compared to WT corresponding cell types. Each bar represents the mean of the relative expression compared to the expression of *GAPDH* in the same sample. Error bars represent SEM of at least three experimental repeats. The RT-qPCR data are presented as mean ± SD from independent triplicates. Statistical analyses were performed using a one-tail two-sample Student's t-test compared to WT (*=p-value < 0.05, **=p-value < 0.01, ***=p value < 0.001).

***Figure S7***

**Expression profile of genes associated with promoter and intragenic hypo-DMRs in ICF1 iPSCs**

(**A**) RT-qPCR of the upregulated *RNF212*, *PTPN20*, *TSPYL5* germline genes in WT, ICF1 and corrected iPSCs, performed as described in Fig. S6E.
(**B**) Left, a genome browser view of *TMEM132D* gene showing intragenic hypo-DMRs in ICF1 iPSCs (highlighted with dashed boxes) and their respective corrected clones. Intragenic CGIs and cis-regulatory

regions (ENCODE) are indicated below in the figure. Right, RT-qPCR of downregulated *TMEM132D* gene in WT, ICF1 and corrected iPSCs, performed as described in Fig. S6E.

(**C**) Left, RT-qPCR analysis of the *VAV1* gene associated with intragenic hypomethylated enhancer-like regions. Each bar represents the mean of the relative expression compared to the expression of *GAPDH* in the same sample. Error bars represent SEM of at least three experimental repeats. Statistical analyses were performed using a one-tail two-sample Student's t-test compared to WT (*=p value < 0.05, **=p value < 0.01, ***=p value < 0.001). Right, a zoom-in genome-browser view of the hypo-DMR associated with the intragenic region (exons 10-13) of *VAV1* gene, as shown in Fig. 3E, which illustrates the methylation coverage in WT1, WT2, pR, cR7 and cR35. Green columns mark CGs hypomethylated in pR iPSCs, showing comparable partial rescue in both corrected clones versus WTs, while the yellow column indicates CGs which remain hypomethylated only in cR7 and not in cR35 compared to WT iPSCs, and contribute to the region being called as a hypo-DMR.

(**D**) RT-qPCR analysis of the *KLHL23* gene associated with intragenic hypomethylated enhancer-like regions.

***Figure S8***

**Hypo-DMRs associated to CGI and GH regulatory elements**

(**A**) Methylation levels of CGIs and 5kb flanking regions in control, ICF1 and corrected iPSCs. CGIs were categorized into tertiles based on their methylation levels (0-1.00) in WT1 iPSCs and the group with the lowest methylation level (0-0.33) is shown. The average methylation level profiles of CGIs, as well as for the +/-5kb flanking regions, are displayed on the right. The left panel represents the corresponding heatmap showing the methylation levels of each hypo-DMR analyzed in the average plots.

(**B**) Distribution of distances between hypo-DMRs in pR and pG iPSCs and CGIs. In each plot the X axis denotes the shortest distance (bp) between the compared regions, while the Y-axis indicates the percentage of hypo-DMRs in each bin size of 2kb.

(**C**) Upset plots displaying the hypo-DMRs at GHs (+/-2kb) in pR (top) and pG (bottom) iPSCs and their corrected counterparts. Vertical bars represent the intersection size between the hypo-DMRs present in each iPSC sample. Black connected dots below each plot represent the hypo-DMRs present in each intersection. Dark red bars correspond to hypo-DMRs present in pR or pG iPSCs (black dot) and absent, therefore rescued, in both cR7 and cR35 (n=4131) or in cG13 and cG50 (n=4139) (grey dots). Pink bars correspond to hypo-DMRs present in pR or pG iPSCs and in one corrected clone, but absent in the second corrected clone. Grey bars represent the hypo-DMRs that are present in patients and corrected iPSCs, and therefore are resistant to *de novo* methylation following the correction of the *DNMT3B* mutations.

(**D**) A genome browser view of the *WDR97* gene, representing an example of methylation loss at enhancer regions (ENCODE cCRE and GH) in pR and pG iPSCs, and ICF1 LCLs. Dark red tracks display methylation coverage measured by WGBS and grey boxes represent hypo-DMRs detected in pR, pG and

corrected iPSCs compared to WT1 and WT2 iPSCs. The tracks underneath indicate the gene regulatory elements including enhancers reported in ENCODE cis Regulatory Elements (cCRE) and in GeneHancer (GH) databases, as well as the H3K4me1, H3K27Ac enrichment and DNAseI sites. Bright red tracks indicate methylation coverage measured by WGBS in ICF1p1 LCLs and WT LCLs and the brown box underneath represents hypo-DMRs in ICF1p1 LCLs.
(**E**) Boxplots representing the distribution of methylation levels following WGBS analysis (expressed as ratio of the number of Cs over the total number of Cs and Ts) at GH promoters and enhancers associated hypo-DMRs in ICF1 iPSCs, which either remain hypomethylated (n=605 for pR and 1024 for pG), or that are rescued (n=4131 for pR and 4139 for pG) in the corresponding isogenic clones.

***Figure S9***

**DNMT3B binding, H3K4me3 and H3K36me3 enrichment at hypo-DMRs**

(**A**) Heatmap representation (left) and average plots (right) of ChIP-seq binding profiles of DNMT3B, H3K36me3 and H3K4me3 in WT1 iPSCs over the expressed genes. The enrichment profiles are sorted with respect to the expression of the genes. Expression levels were determined by RNA-seq, calculated as log2 of FPKM counts in WT1. The highest levels of expression are at the bottom, the lowest ones are at the top. For plots on the right, DNMT3B and histone mark enrichments at expressed genes in WT1 are clustered into four quartiles based on gene expression levels (blue lines; Q1 – lowest expressed genes, Q4 – highest expressed genes). For DNMT3B and H3K36me3, the X-axis denotes genomic regions spanning +/-2kb of gene bodies. For H3K4me3 the X-axis denotes genomic regions spanning +/-2kb of TSSs. The Y-axis for each ChIP-Seq dataset indicates the number of counts per million mapped reads within regions of expressed genes.
(**B**) Distribution of distances between hypo-DMRs and increased H3K4me3 DERs in pR (left) and pG (right) iPSCs compared to WT1. In each plot the X-axis denotes the shortest distance (bp) between the compared regions, while the Y-axis indicates the percentage of hypo-DMRs in each bin of 1kb size.
(**C**) Plots of average CG methylation levels (expressed as ratio of the number of Cs over the total number of Cs and Ts) at increased H3K4me3 DERs in pR (n=2,823; left) and pG (n=1,178; right) iPSCs in comparison to the control WT1 and WT2 iPSCs.
(**D**) Plots of average CG methylation levels of hypo-DMRs intersecting with increased H3K4me3 DERs (+/-2kb) in pR (n=1,442, left) and in pG (n=1,081, right) in comparison to the control WT1 and WT2 iPSCs.
(**E**) ChIP-qPCR measuring H3K4me3 levels at *PCDHGA6* and *TNXB* genes in WT, ICF1 and corrected iPSCs. Amplicon enrichment in immunoprecipitated and mock samples (orange bars) is expressed as percentage (%) of input. Bars and error bars represent means and SEM of at least three experimental repeats. Statistical analyses were performed using a one-tail two-sample Student's t-test compared to WT1 (**=p value < 0.01, ***=p value < 0.001). Primer sets used for ChIP analysis are reported in Table S2.

**Table S1. Sequencing experimental design**

| Experiment | Sequencing reads | Read length | Number of sequenced reads per sample |
|---|---|---|---|
| **WGBS** | Paired-end (PE) | 100bp | 300-350 million |
| **ChIP-seq**<br>**i) H3K4me3**<br>**ii) H3K36me3**<br>**iii) DNMT3B** | Single-end (SE) | 100bp | <br>25-30 million<br>40-50 million<br>50-60 million |
| **RNA-seq** | Paired-end (PE) | 125bp | 50-60 million |

**Table S2. List of primer sequences and qPCR conditions used in RTqPCR experiments**

| Gene | Primer Sequence 5'→ 3' | Experiment; amplicon size (bp) | Thermocycling parameters |
|---|---|---|---|
| *RUNX1* | F- CCACCTACCACAGAGCCATCAA (exon 5 ENST00000300305.7)<br>R- TTCACTGAGCCGCTCGGAAAAG (exon 6 ENST00000300305.7) | RT-qPCR; 139 bp | 95°C-62°C-72°C<br>20" -20" -20" (35 cycles) |
| *IGHG1* | F- CGGATATGGCTCTTGGCAGG (intron 1 ENST00000390548.6)<br>R- TTCTGGCTTTTTCCCCAGGC (intron 1 ENST00000390548.6) | RT-qPCR; 120 bp | 95°C-62°C-72°C<br>20" -20" -20" (35 cycles) |
| *LINC00221* | F- GGTAGCTCAGCGGGGACTT (exon 1 ENST00000619530.1)<br>R- CTCTCCCAGCCAGGACCTC (exon 1 ENST00000619530.1) | RT-qPCR; 90 bp | 95°C-62°C-72°C<br>20" -20" -20" (35 cycles) |
| *CD74* | F- AGGCACAGGGAGAAGGGATA (3'UTR ENST00000009530.13)<br>R- GGCTCAGCCTGCAGGTAAAT (3'UTR ENST00000009530.13) | RT-qPCR; 117 bp | 95°C-62°C-72°C<br>20" -20" -20" (35 cycles) |
| *PLA2G4C* | F- GACAAGATAATGAGCAGCCGGAAG (exon 14 ENSG00000105499.14) | RT-qPCR; 129 bp | 95°C-62°C-72°C<br>20" -20" -20" (35 cycles) |

| | | | |
|---|---|---|---|
| | R- GGCACTGAAGTCGAAGGAGAGG<br>(exon 14 ENSG00000105499.14) | | |
| *RNF212* | F- TGCTTGATTTGTAAAGCTCCTTG<br>(exon 2 ENSG00000178222.13)<br>R- TGGGAGGTTTCCCTGGAGTA<br>(exon 2 ENSG00000178222.13) | RT-qPCR; 141 bp | 95°C-62°C-72°C<br>20” -20” -20” (35 cycles) |
| *PTPN20* | F- CCTGTTGGTCTGGGAAGCAT<br>(3’UTR ENST00000374339.5)<br>R- AGGCATGGCAAAAGTCTCCT<br>(3’UTR ENST00000374339.5) | RT-qPCR; 148 bp | 95°C-62°C-72°C<br>20” -20” -20” (35 cycles) |
| *TSPYL5* | F- CGTGTCTTTGAAGCTGCCTCC<br>(exon 1 ENST00000322128.5)<br>R- TACTGTGAAGGGTCCGGGTC<br>(exon 1 ENST00000322128.5) | RT-qPCR; 152 bp | 95°C-64°C-72°C<br>20” -20” -20” (35 cycles) |
| *TMEM132D* | F- TGCAGCCAGTAAGTCACCTG<br>(3’UTR ENST00000422113.7)<br>R- CGCAGATGGATTTGGAAGCG<br>(3’UTR ENST00000422113.7) | RT-qPCR; 188 bp | 95°C-62°C-72°C<br>20” -20” -20” (35 cycles) |
| *VAV1* | F- CACCTTGCAGTTCCCCTTCA<br>(exon 25 ENST00000602142.6)<br>R- GACAGCTCTGATCGGTCTCG<br>(exon 26 ENST00000602142.6) | RT-qPCR; 132 bp | 95°C-62°C-72°C<br>20” -20” -20” (35 cycles) |
| *KLHL23* | F- GAAAGGAGGATGGAGTGCGG<br>(exon 4 ENST00000272797.8)<br>R- CAAACACACCCATGAGACCG<br>(exon 4 ENST00000272797.8) | RT-qPCR; 176 bp | 95°C-62°C-72°C<br>20” -20” -20” (35 cycles) |
| *GAPDH* | F- GAAGGTGAAGGTCGGAGTC<br>(exon 2 ENST00000229239.10)<br>R- GAAGATGGTGATGGGATTTC<br>(exon 3 ENST00000229239.10) | Normalizing gene for RT-qPCR; 234 bp | 95°C-62°C or 64°C-72°C<br>20”-20”-20” (35 cycles) |

| | | | |
|---|---|---|---|
| *PCDHGA6* | F- TAAGCCAGTAATGGCGCCTC<br>(exon 1 ENSG00000253731.3)<br>R- CCAGTCCCAGATCCTTGACG<br>(exon 1 ENSG00000253731.3) | ChIP-qPCR; 172 bp | 95°C-62°C-72°C<br>20” -20” -20” (39 cycles) |
| *TNXB* | F- ATTTGCGACACGGGCTACAG<br>(exon 1 ENSG00000168477.19)<br>R- AAACACACACGCCGTTCTC<br>(exon 1 ENSG00000168477.19) | ChIP-qPCR; 190 bp | 95°C-64°C-72°C<br>20” -20” -20” (39 cycles) |
| *MYOD1* | F- CCTCTTTCGGTCCCTCTTTC<br>(promoter ENSG00000129152.4)<br>R- TTCCAAACCTCTCCAACACC<br>(promoter ENSG00000129152.4) | Assessing the absence of genomic DNA in RT-qPCR; 223 bp | 95°C-62°C-72°C<br>20” -20” -20” (39 cycles) |

**Table S3**

Lists of hypo-DMRs identified in ICF1 iPSCs and their categorization based on genomic annotation, rescue status in isogenic corrected clones and overlap with CGI and/or GeneHancer (GH) regulatory elements. Hypo-DMR score denotes the difference in methylation percentages between the ICF1 and WT1 iPSCs. Group assignment and rescue category (full, partial, no rescue) of each hypo-DMR is reported in Fig. 1A.

List of biological processes (PANTHER) enriched among the genes associated with hypo-DMRs identified in ICF1 iPSCs and annotated to promoter or gene body. The column IDs are indicated as provided by the PANTHER software.

HOMER results showing the known Transcription Factor Motifs enriched at hypo-DMRs identified in ICF1 iPSCs and overlapping with decreased DNMT3B DERs (DNMT3B-Dec).

**Table S4**

Genes that are differentially expressed (DE; pp > 0.9) and genes with pp > 0.8 obtained from RNA-Seq data analysis of ICF1 iPSCs compared to WT1. Log2FC and posterior probability for both ICF1 and the corresponding corrected clones compared with WT1 are provided for the listed genes.

**Table S5**

HOMER results showing the known Transcription Factor Motifs enriched at the subset of hypo-DMRs belonging to Group1 and 2 or Group 3 and 4, based on the definition described in Fig. 1A. The column IDs are indicated as provided by the HOMER software.

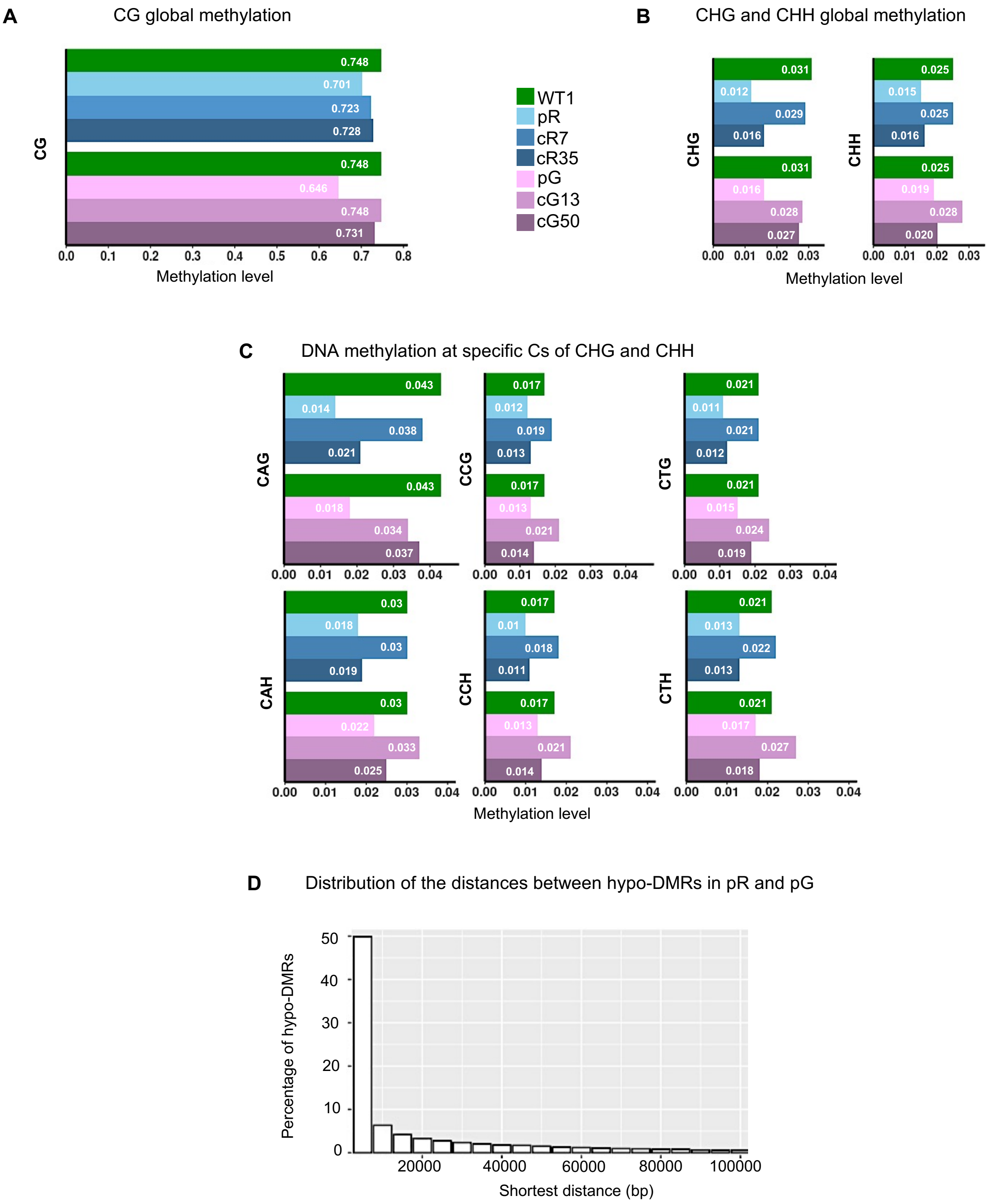
A
CG global methylation
CG
0.748
0.701
0.723
0.728
0.748
0.646
0.748
0.731
0.0 0.1 0.2 0.3 0.4 0.5 0.6 0.7 0.8
Methylation level
WT1
pR
cR7
cR35
pG
cG13
cG50
B
CHG and CHH global methylation
CHG
0.031
0.012
0.029
0.016
0.031
0.016
0.028
0.027
CHH
0.025
0.015
0.025
0.016
0.025
0.019
0.028
0.020
0.00 0.01 0.02 0.03
Methylation level
C
DNA methylation at specific Cs of CHG and CHH
CAG
0.043
0.014
0.038
0.021
0.043
0.018
0.034
0.037
CCG
0.017
0.012
0.019
0.013
0.017
0.013
0.021
0.014
CTG
0.021
0.011
0.021
0.012
0.021
0.015
0.024
0.019
CAH
0.03
0.018
0.03
0.019
0.03
0.022
0.033
0.025
CCH
0.017
0.01
0.018
0.011
0.017
0.013
0.021
0.014
CTH
0.021
0.013
0.022
0.013
0.021
0.017
0.027
0.018
0.00 0.01 0.02 0.03 0.04
Methylation level
D
Distribution of the distances between hypo-DMRs in pR and pG
Percentage of hypo-DMRs
0 10 20 30 40 50
20000 40000 60000 80000 100000
Shortest distance (bp)

*Figure S1*

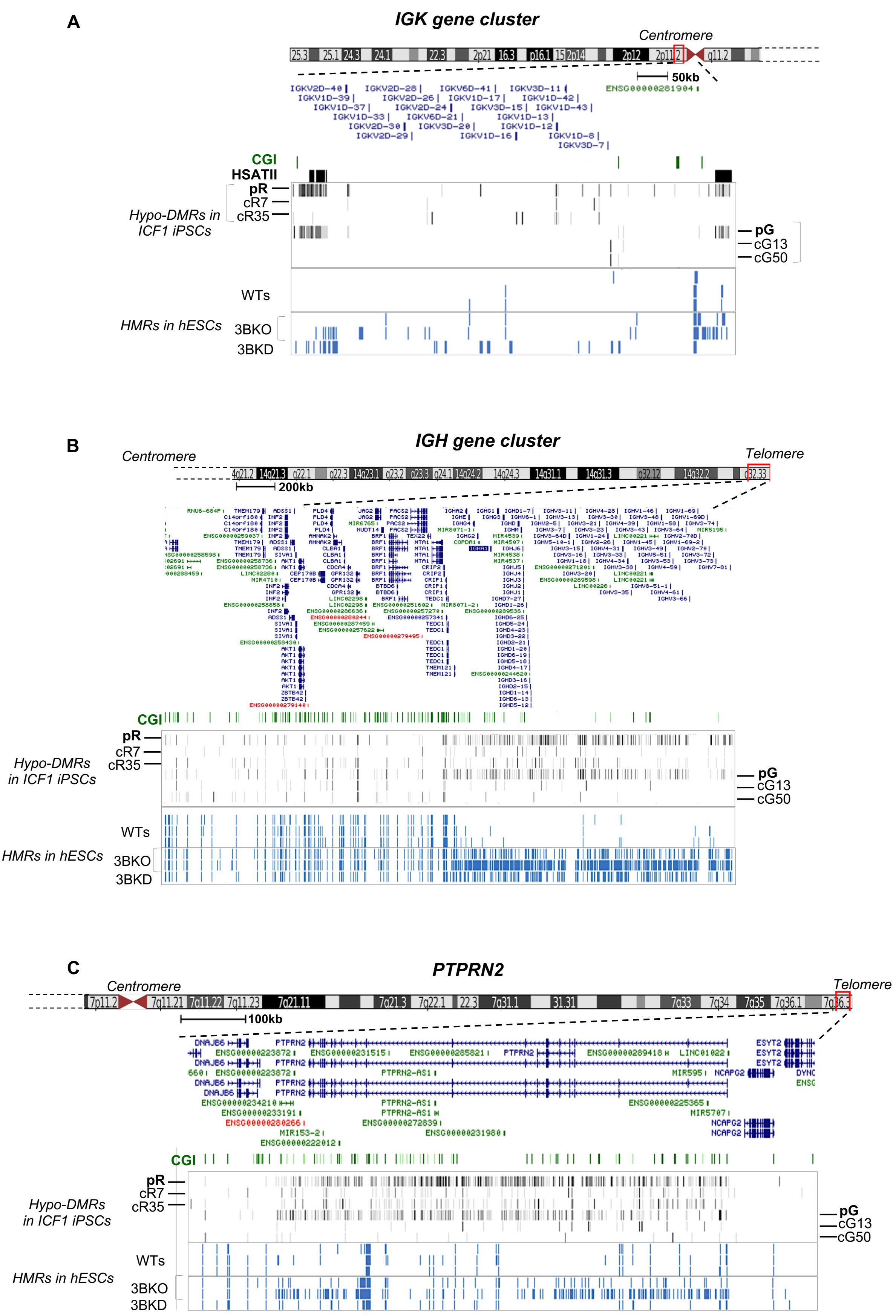
A
IGK gene cluster
Centromere
50kb
CGI
HSATII
pR
cR7
cR35
Hypo-DMRs in ICF1 iPSCs
pG
cG13
cG50
WTs
HMRs in hESCs
3BKO
3BKD
B
IGH gene cluster
Centromere
Telomere
200kb
CGI
pR
cR7
cR35
Hypo-DMRs in ICF1 iPSCs
pG
cG13
cG50
WTs
HMRs in hESCs
3BKO
3BKD
C
PTPRN2
Centromere
Telomere
100kb
CGI
pR
cR7
cR35
Hypo-DMRs in ICF1 iPSCs
pG
cG13
cG50
WTs
HMRs in hESCs
3BKO
3BKD

*Figure S2*

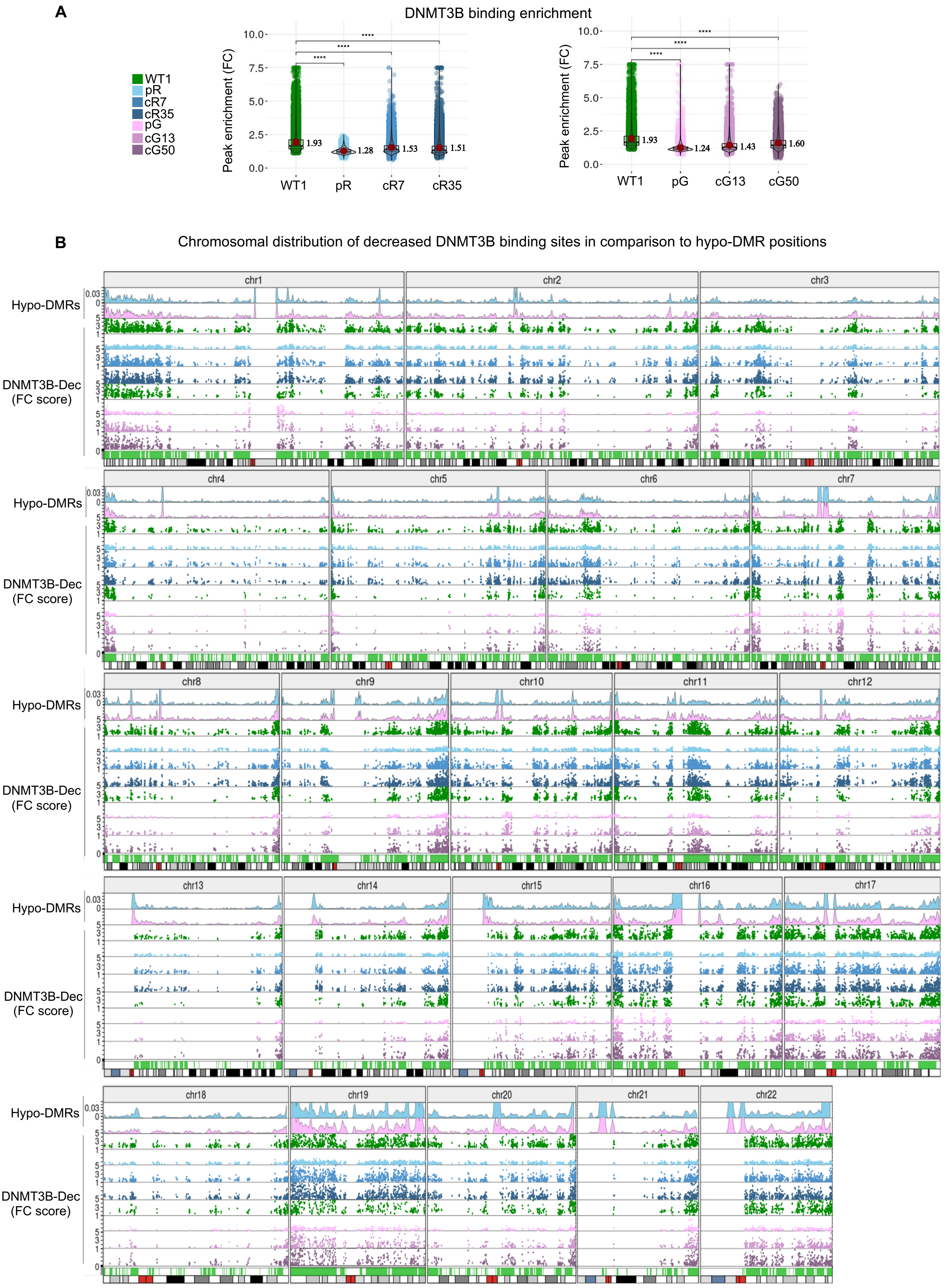
A
DNMT3B binding enrichment
WT1
pR
cR7
cR35
pG
cG13
cG50
Peak enrichment (FC)
10.0
7.5
5.0
2.5
0.0
****
1.93
1.28
1.53
1.51
1.24
1.43
1.60
B
Chromosomal distribution of decreased DNMT3B binding sites in comparison to hypo-DMR positions
Hypo-DMRs
DNMT3B-Dec
(FC score)
chr1
chr2
chr3
chr4
chr5
chr6
chr7
chr8
chr9
chr10
chr11
chr12
chr13
chr14
chr15
chr16
chr17
chr18
chr19
chr20
chr21
chr22

*Figure S3*

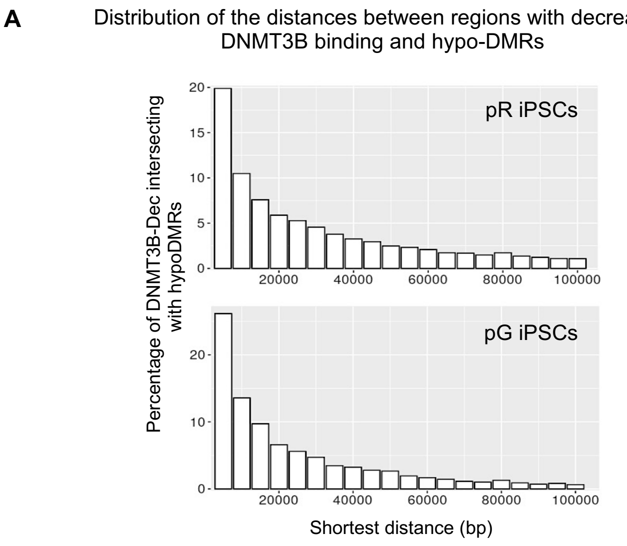
A
Distribution of the distances between regions with decreased DNMT3B binding and hypo-DMRs
Percentage of DNMT3B-Dec intersecting with hypoDMRs
pR iPSCs
pG iPSCs
Shortest distance (bp)

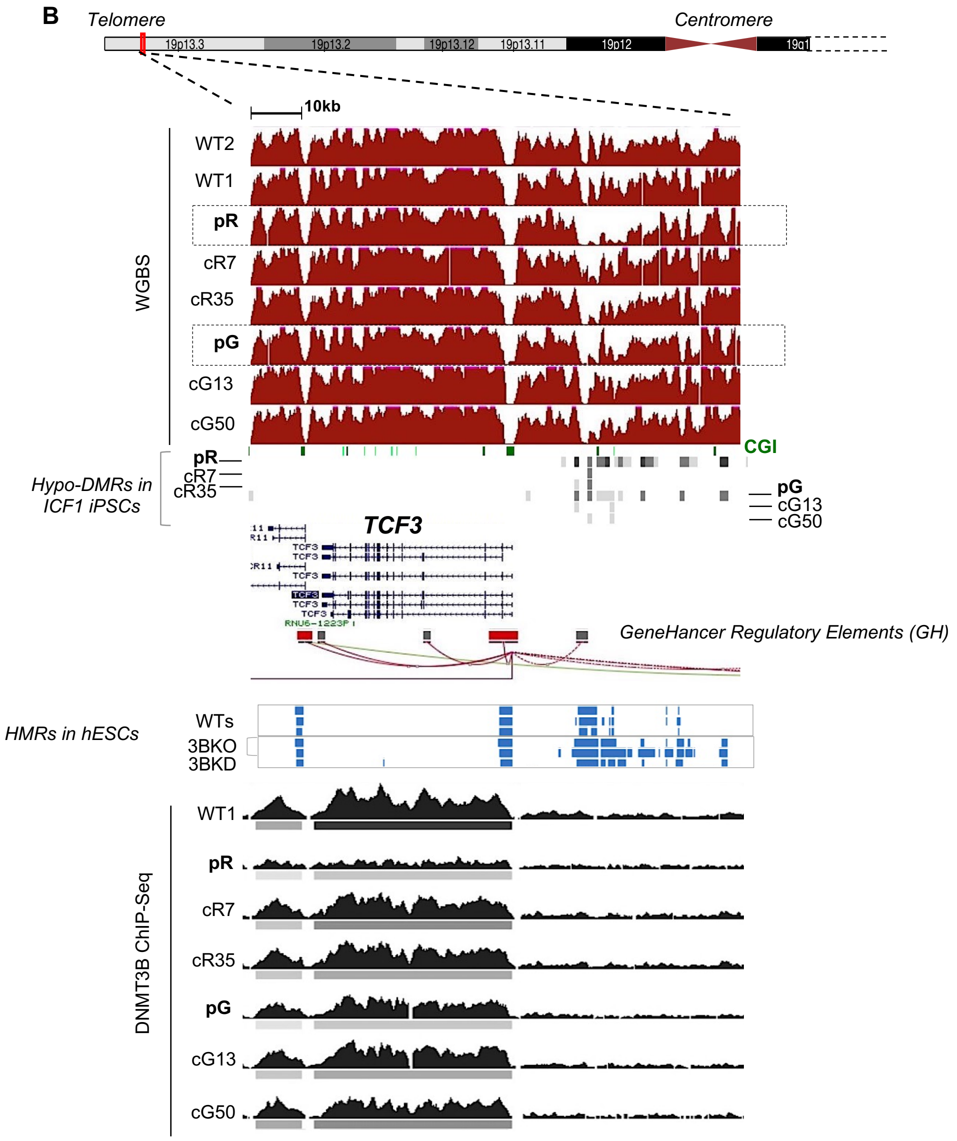
B
Telomere
Centromere
19p13.3
19p13.2
19p13.12
19p13.11
19p12
10kb
WGBS
WT2
WT1
pR
cR7
cR35
pG
cG13
cG50
CGI
Hypo-DMRs in ICF1 iPSCs
pR
cR7
cR35
pG
cG13
cG50
TCF3
GeneHancer Regulatory Elements (GH)
HMRs in hESCs
WTs
3BKO
3BKD
DNMT3B ChIP-Seq
WT1
pR
cR7
cR35
pG
cG13
cG50

**Figure S4**

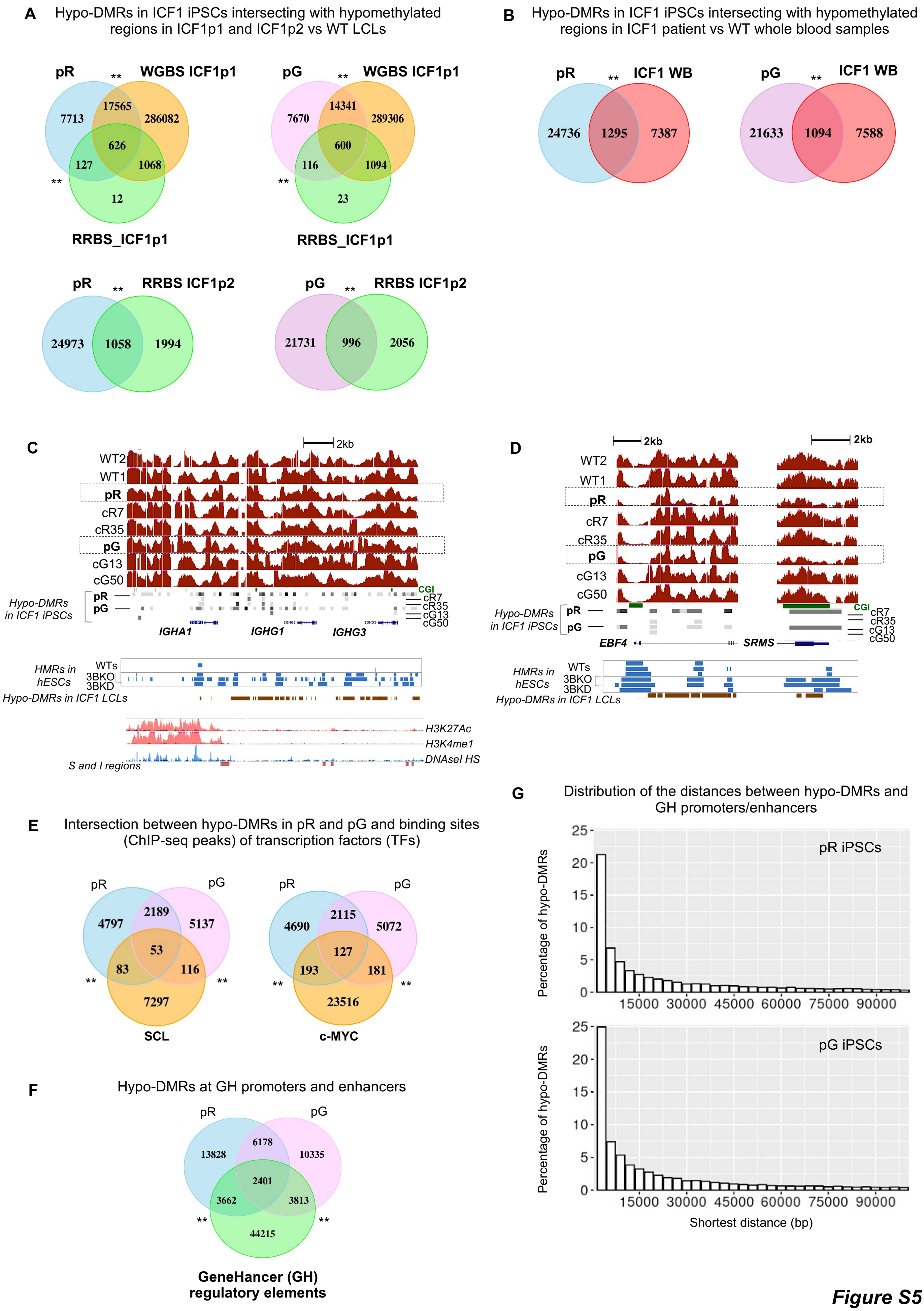
A
Hypo-DMRs in ICF1 iPSCs intersecting with hypomethylated regions in ICF1p1 and ICF1p2 vs WT LCLs
pR
WGBS ICF1p1
RRBS_ICF1p1
7713
17565
286082
626
127
1068
12
pG
7670
14341
289306
600
116
1094
23
RRBS ICF1p2
24973
1058
1994
21731
996
2056
B
Hypo-DMRs in ICF1 iPSCs intersecting with hypomethylated regions in ICF1 patient vs WT whole blood samples
ICF1 WB
24736
1295
7387
21633
1094
7588
C
2kb
WT2
WT1
pR
cR7
cR35
pG
cG13
cG50
Hypo-DMRs in ICF1 iPSCs
CGI
IGHA1
IGHG1
IGHG3
HMRs in hESCs
WTs
3BKO
3BKD
Hypo-DMRs in ICF1 LCLs
H3K27Ac
H3K4me1
DNAseI HS
S and I regions
D
2kb
EBF4
SRMS
E
Intersection between hypo-DMRs in pR and pG and binding sites (ChIP-seq peaks) of transcription factors (TFs)
4797
2189
5137
53
83
116
7297
SCL
4690
2115
5072
127
193
181
23516
c-MYC
F
Hypo-DMRs at GH promoters and enhancers
13828
6178
10335
2401
3662
3813
44215
GeneHancer (GH) regulatory elements
G
Distribution of the distances between hypo-DMRs and GH promoters/enhancers
Percentage of hypo-DMRs
pR iPSCs
pG iPSCs
Shortest distance (bp)
0
5
10
15
20
25
15000
30000
45000
60000
75000
90000

*Figure S5*

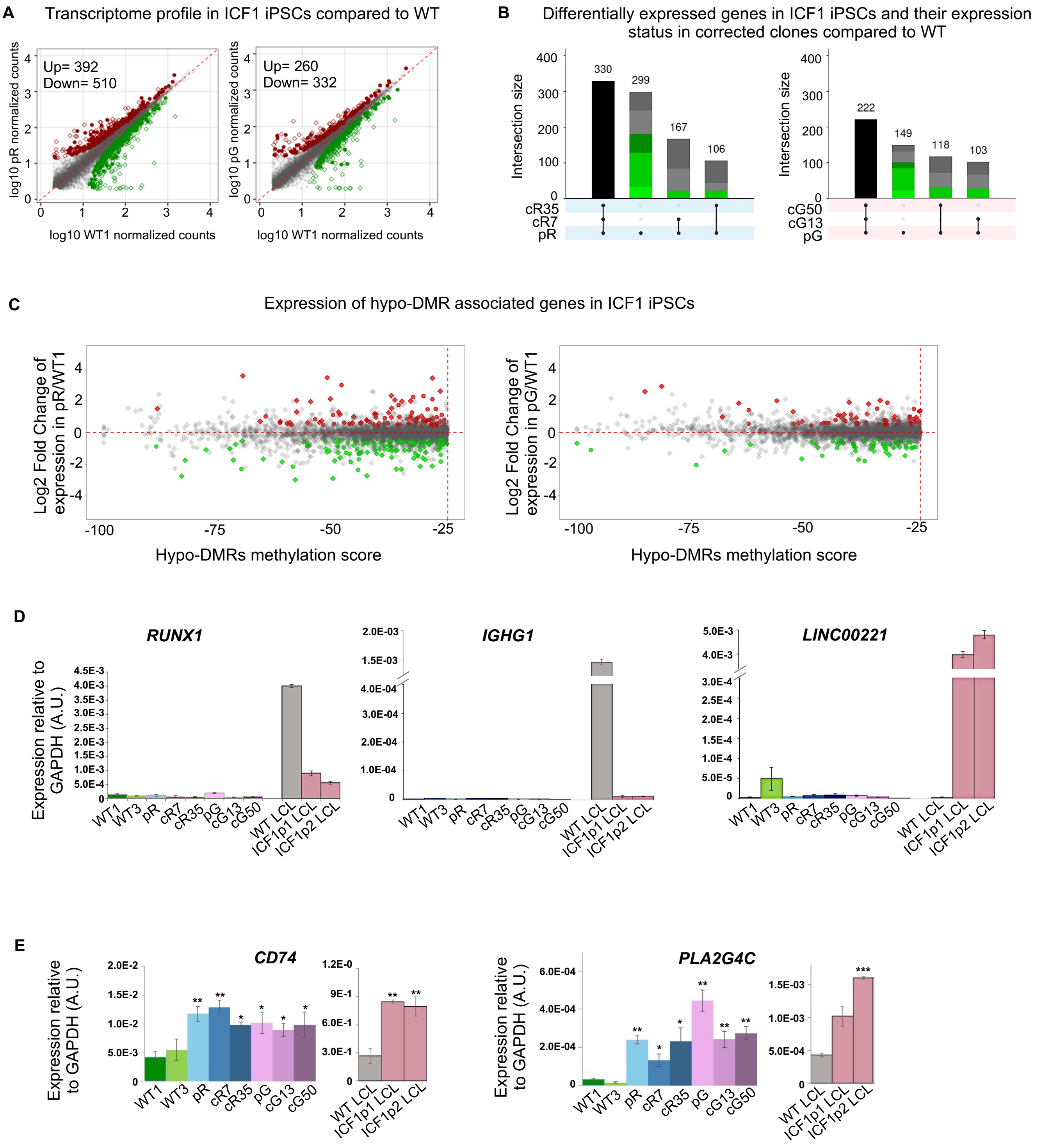
A
Transcriptome profile in ICF1 iPSCs compared to WT
Up= 392
Down= 510
Up= 260
Down= 332
log10 pR normalized counts
log10 pG normalized counts
log10 WT1 normalized counts
B
Differentially expressed genes in ICF1 iPSCs and their expression status in corrected clones compared to WT
Intersection size
330
299
167
106
222
149
118
103
cR35
cR7
pR
cG50
cG13
pG
C
Expression of hypo-DMR associated genes in ICF1 iPSCs
Log2 Fold Change of expression in pR/WT1
Log2 Fold Change of expression in pG/WT1
Hypo-DMRs methylation score
D
RUNX1
IGHG1
LINC00221
Expression relative to GAPDH (A.U.)
WT1
WT3
pR
cR7
cR35
pG
cG13
cG50
WT LCL
ICF1p1 LCL
ICF1p2 LCL
E
CD74
PLA2G4C
Expression relative to GAPDH (A.U.)

*Figure S6*

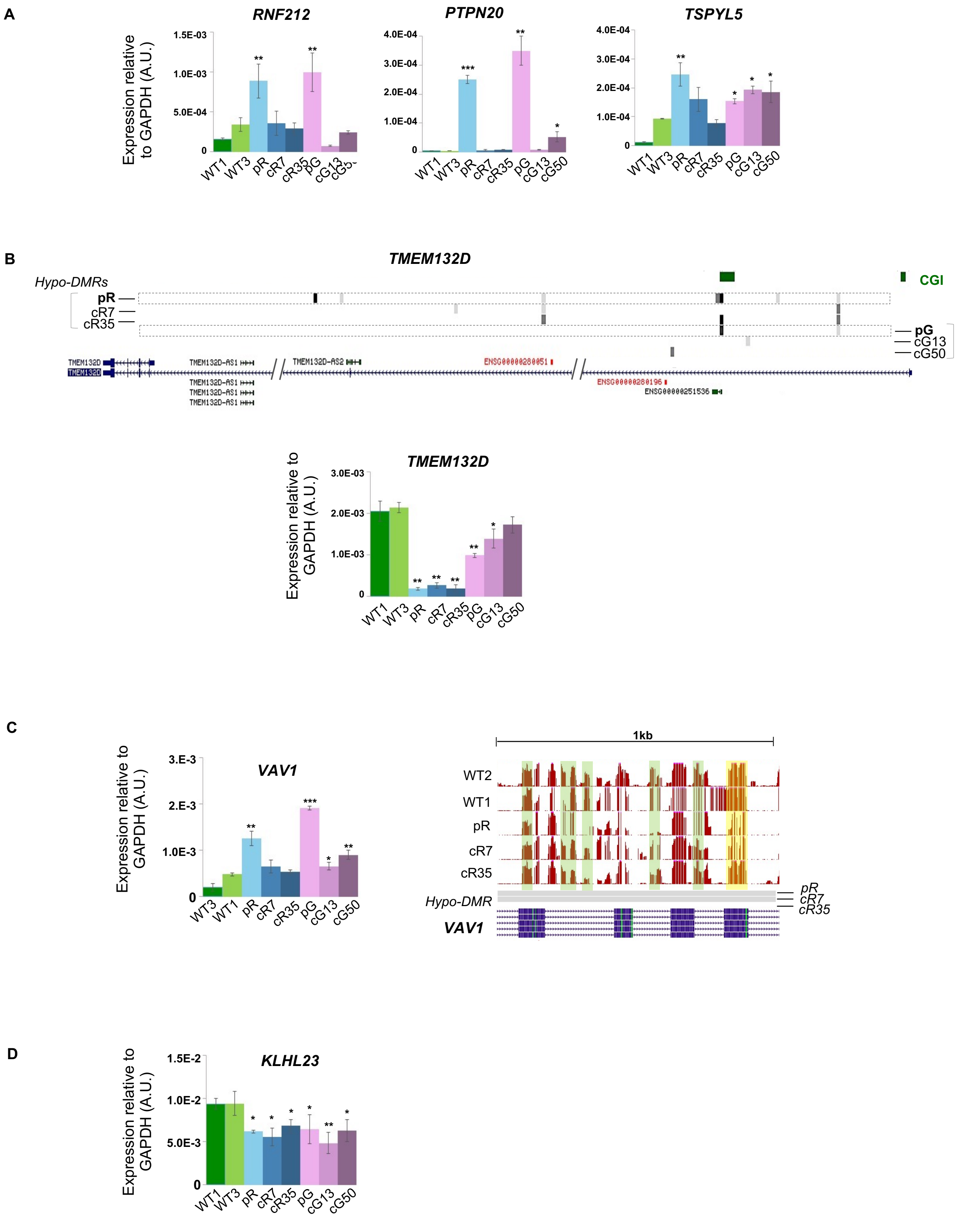
A
RNF212
PTPN20
TSPYL5
Expression relative to GAPDH (A.U.)
1.5E-03
1.0E-03
5.0E-04
0
4.0E-04
3.0E-04
2.0E-04
1.0E-04
WT1 WT3 pR cR7 cR35 pG cG13 cG50
B
TMEM132D
Hypo-DMRs
pR
cR7
cR35
pG
cG13
cG50
CGI
TMEM132D-AS1
TMEM132D-AS2
ENSG00000280051
ENSG00000280196
ENSG00000251536
3.0E-03
2.0E-03
1.0E-03
C
VAV1
3.E-3
2.E-3
1.0E-3
1kb
WT2
WT1
pR
cR7
cR35
Hypo-DMR
D
KLHL23
1.5E-2
1.0E-2
5.0E-3

**Figure S7**

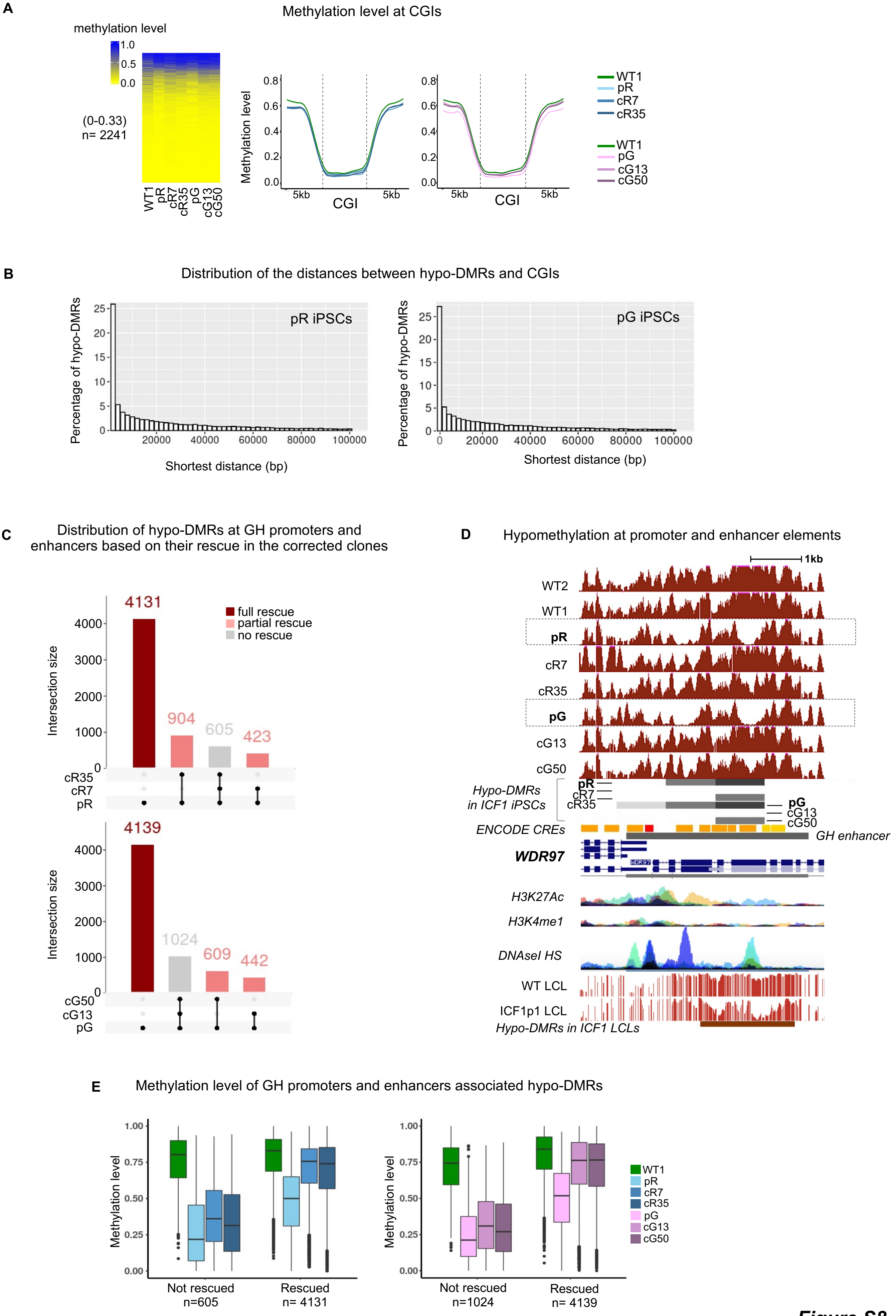
A
Methylation level at CGIs
methylation level
1.0
0.5
0.0
(0-0.33)
n= 2241
WT1
pR
cR7
cR35
pG
cG13
cG50
Methylation level
0.8
0.6
0.4
0.2
0.0
5kb
CGI
5kb
WT1
pR
cR7
cR35
WT1
pG
cG13
cG50
B
Distribution of the distances between hypo-DMRs and CGIs
Percentage of hypo-DMRs
pR iPSCs
pG iPSCs
25
20
15
10
5
0
20000
40000
60000
80000
100000
Shortest distance (bp)
C
Distribution of hypo-DMRs at GH promoters and enhancers based on their rescue in the corrected clones
Intersection size
4131
904
605
423
full rescue
partial rescue
no rescue
4000
3000
2000
1000
0
cR35
cR7
pR
4139
1024
609
442
cG50
cG13
pG
D
Hypomethylation at promoter and enhancer elements
1kb
WT2
WT1
pR
cR7
cR35
pG
cG13
cG50
Hypo-DMRs in ICF1 iPSCs
pR
cR7
cR35
pG
cG13
cG50
ENCODE CREs
GH enhancer
WDR97
H3K27Ac
H3K4me1
DNAseI HS
WT LCL
ICF1p1 LCL
Hypo-DMRs in ICF1 LCLs
E
Methylation level of GH promoters and enhancers associated hypo-DMRs
Methylation level
1.00
0.75
0.50
0.25
0.00
Not rescued
n=605
Rescued
n= 4131
Not rescued
n=1024
Rescued
n= 4139
WT1
pR
cR7
cR35
pG
cG13
cG50

***Figure S8***

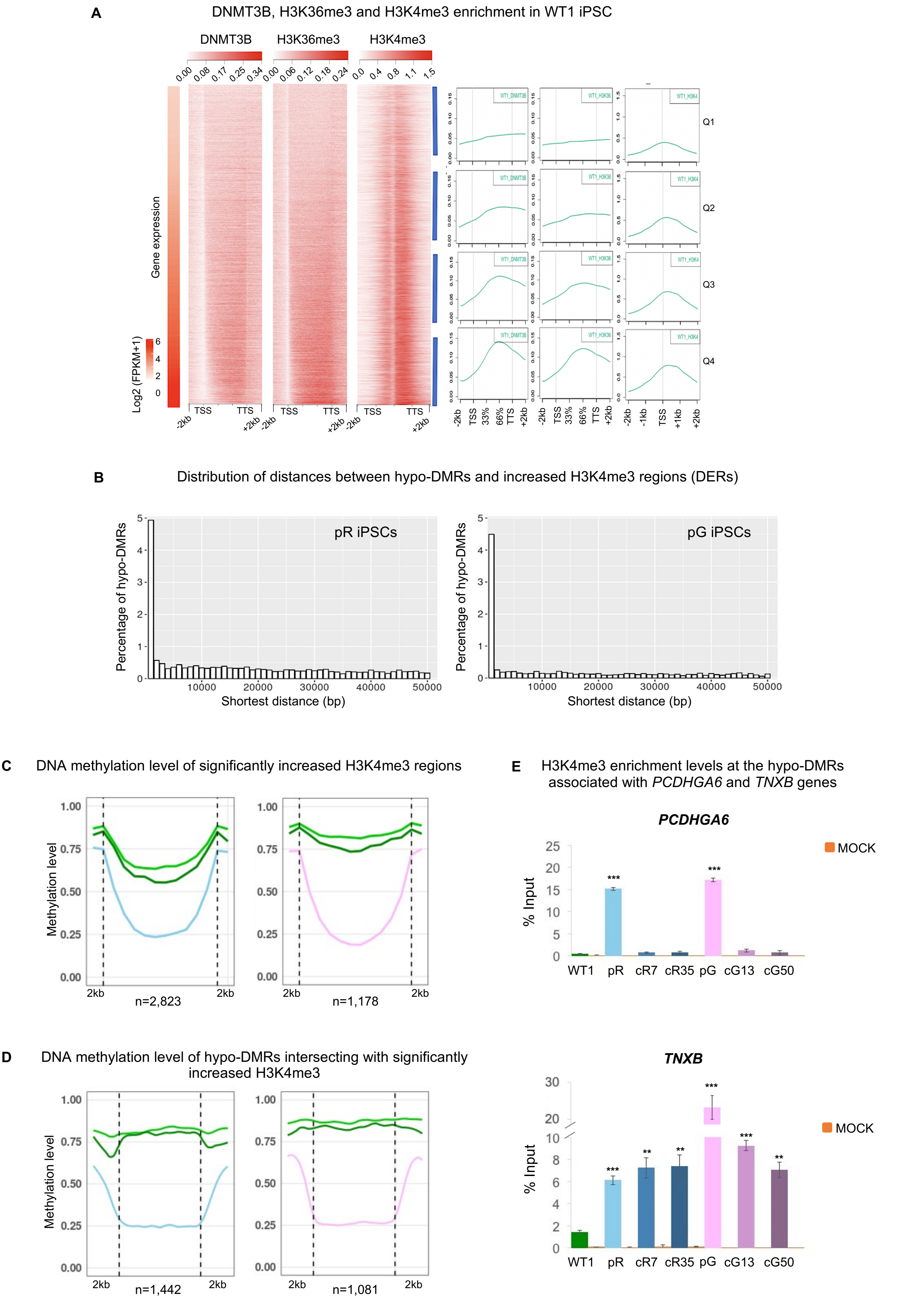
A
DNMT3B, H3K36me3 and H3K4me3 enrichment in WT1 iPSC
DNMT3B
H3K36me3
H3K4me3
Gene expression
Log2 (FPKM+1)
TSS
TTS
-2kb
+2kb
Q1
Q2
Q3
Q4
B
Distribution of distances between hypo-DMRs and increased H3K4me3 regions (DERs)
pR iPSCs
pG iPSCs
Percentage of hypo-DMRs
Shortest distance (bp)
C
DNA methylation level of significantly increased H3K4me3 regions
Methylation level
2kb
n=2,823
n=1,178
D
DNA methylation level of hypo-DMRs intersecting with significantly increased H3K4me3
n=1,442
n=1,081
E
H3K4me3 enrichment levels at the hypo-DMRs associated with PCDHGA6 and TNXB genes
PCDHGA6
TNXB
% Input
MOCK
WT1
pR
cR7
cR35
pG
cG13
cG50

*Figure S9*
